# Compartmentalization of the replication fork by single-stranded DNA binding protein regulates translesion synthesis

**DOI:** 10.1101/2020.03.03.975086

**Authors:** Seungwoo Chang, Elizabeth S. Thrall, Luisa Laureti, Vincent Pagès, Joseph J. Loparo

**Author notes:** Department of Chemistry, Fordham University, Bronx, NY.

## Abstract

DNA replication is mediated by the coordinated actions of multiple enzymes within replisomes. Processivity clamps tether many of these enzymes to DNA, allowing access to the primer/template junction. Many clamp-interacting proteins (CLIPs) are involved in genome maintenance pathways including translesion synthesis (TLS). Despite their abundance, DNA replication in bacteria is not perturbed by these CLIPs. Here we show that while the TLS polymerase Pol IV is largely excluded from moving replisomes, the remodeling of ssDNA binding protein (SSB) upon replisome stalling enriches Pol IV at replication forks. This enrichment is indispensable for Pol IV-mediated TLS on both the leading and lagging strands as it enables Pol IV-processivity clamp binding by overcoming the gatekeeping role of the Pol III epsilon subunit. As we have demonstrated for the Pol IV-SSB interaction, we propose that the binding of CLIPs to the processivity clamp must be preceded by interactions with factors that serve as localization markers for their site of action.

## Introduction

DNA damage is a potent challenge to the replisome. Cells employ a vast array of enzymatic activities to either tolerate or repair this damage and enable DNA replication. However, inappropriate use of these enzymes can contribute to genome instability. During translesion synthesis (TLS), a prominent damage tolerance pathway, error-prone polymerases must gain access to the primer template (P/T) junction to extend the nascent strand past a blocking DNA lesion. Given their low fidelity, access must be tightly regulated to ensure that TLS polymerases are used only when necessary.

Polymerases and other repair factors are often tethered to their DNA substrates through interaction with processivity clamps. These clamps are multimeric, ring-shaped molecules that encircle DNA and interact with clamp interacting proteins (“CLIPs”) through conserved binding surfaces. Within *E. coli* all 5 DNA polymerases interact with the β_2_ clamp, along with a number of other factors involved in Okazaki fragment maturation (LigA) (López de Saro and O’Donnell, 2001), regulation of replication initiation (Hda) (Kurz et al., 2004), genome arrangement (CrfC) (Ozaki et al., 2013) and mismatch repair (MutS and MutL) (López de Saro and O’Donnell, 2001). CLIPs contain one or more clamp binding motifs (CBMs), which mediate interaction with the β_2_ clamp. These motifs are required for CLIP function in the cell and ensure their actions at proper sites. Cellular copy numbers of CLIPs and their binding affinities to the β_2_ clamp vary vastly, and the copy numbers of TLS polymerases increase during the SOS DNA damage response.

How a large pool of CLIPs competes for a limited number of clamp binding sites remains unclear. CLIP occupancy on the clamp may be largely determined by the relative abundance of CLIPs and their binding affinities to the clamp. However, this model does not clearly explain how highly abundant CLIPs, such as TLS polymerases, do not prevent the clamp binding of less abundant CLIPs. For example, processive replication mediated by Pol III, whose copy number is around 20, is only marginally inhibited by TLS polymerases, whose combined copy number is 1-2 orders of magnitude greater than that of Pol III depending on SOS induction(Tan et al., 2015). Intriguingly, most CLIPs interact with other factors and/or specific DNA structures that may facilitate clamp binding(Bhattacharyya et al., 2014). Supporting this idea, we previously observed that disrupting the interaction of the TLS polymerase Pol IV with the β_2_ clamp only partially reduced enrichment of Pol IV near stalled replisomes in cells(Thrall et al., 2017), suggesting that Pol IV uses distinct molecular interactions for localization to its site of action and execution of its biochemical functions. Such binary interactions can actively enrich a subset of CLIPs at a specific site, such as replication forks, while passively exclude others.

A subset of CLIPs interacts with ssDNA binding protein (SSB) including all three TLS polymerases(Arad et al., 2008; Furukohri et al., 2012; Molineux and Gefter, 1974). In cells, SSB rapidly associates with ssDNA to protect it from nucleolytic cleavage and chemical damage. Additionally, SSB promotes genome maintenance processes by interacting with a host of cellular factors (SSB Interacting Proteins “SIPs”). *E. coli* SSB serves as an important model system, retaining many of the structural and regulatory features of SSBs including a highly conserved C-terminal peptide (EcSSB-Ct, MDFDDDIPF) that interacts with SIPs.

In this study we demonstrate that Pol IV is absolutely required to interact with replisome-associated SSB to carry out TLS at the replication fork. This interaction enriches Pol IV at lesion-stalled replication forks. The resulting increase in local Pol IV concentration allows for it to overcome a gatekeeping kinetic barrier imposed by the ε subunit of the Pol III core complex(Chang et al., 2019), which competitively inhibits association of Pol IV and other factors with the β_2_ clamp. Interestingly, SSB promotes Pol IV-mediated TLS on both the leading and lagging strands, suggesting a similar gatekeeping mechanism operates on each strand. By promoting TLS, the Pol IV-SSB interaction suppresses resolution of lesion-stalled replisomes through the recombination-dependent damage avoidance pathway, revealing a critical role of SSB in resolution pathway choice and thus damage-induced mutagenesis.

## Results

### Pol IV directly interacts with the SSB through the C-terminal peptide

In an effort to better characterize the Pol IV-SSB interaction, we first asked whether Pol IV interacts with the C-terminal peptide of SSB (SSB-Ct), a conserved binding motif that mediates interaction with other SIPs(Lu and Keck, 2008; Marceau et al., 2011; Shereda et al., 2008; 2009). Consistent with Pol IV directly interacting with SSB, addition of increasing amounts of purified Pol IV to a fluorescein-labeled SSB C-terminal peptide (FL-SSB-Ct) led to a gradual increase in fluorescence polarization (FP) of FL (Fig 1A and 1B). To determine if the Pol IV-SSB-Ct interaction retains common binding features of other SIPs, we next examined how unlabeled SSB-Ct variants competed with FL-SSB-Ct. Increasing amounts of unlabeled SSB-Ct variants (competitor) readily displaced bound FL-SSB-Ct (tracer peptide) from Pol IV as measured by FP (Fig 1C), indicating that the labeled and unlabeled peptides bind the same interaction surface. Furthermore, deletion of the absolutely conserved ultimate phenylalanine (SSB-Ct^ΔF^), a critical interaction residue for other SIPs, substantially reduced binding affinity (Fig. 1C). Mutating a cluster of aspartic acids in the SSB-Ct to alanine (SSB-Ct^ΔD^), which weakens electrostatic interactions of the SSB-Ct with basic ridges found in other SIPs, also reduced binding affinity but less severely than the deletion of the ultimate phenylalanine (Fig. 1C). Combining these two mutations (SSB-Ct^ΔD,ΔF^) completely abolished binding (Fig. 1C). Collectively, these results indicate that the SSB-Ct-binding surface(s) within Pol IV retains structural features found in SSB-Ct-binding surfaces of other SIPs (Fig S1A and B). However, as the cellular concentrations of SSB_4_ (∼500 nM, equivalent to ∼2 uM SSB-Ct) and Pol IV (∼200/∼2000 nM for uninduced/fully induced) are more than 10 times lower than the K_D_ of SSB-Ct (30 μM), binding of Pol IV to the tetrameric complex of full-length SSB (SSB_4_) might be tighter than that to the isolated SSB-Ct.

**Figure 1.**
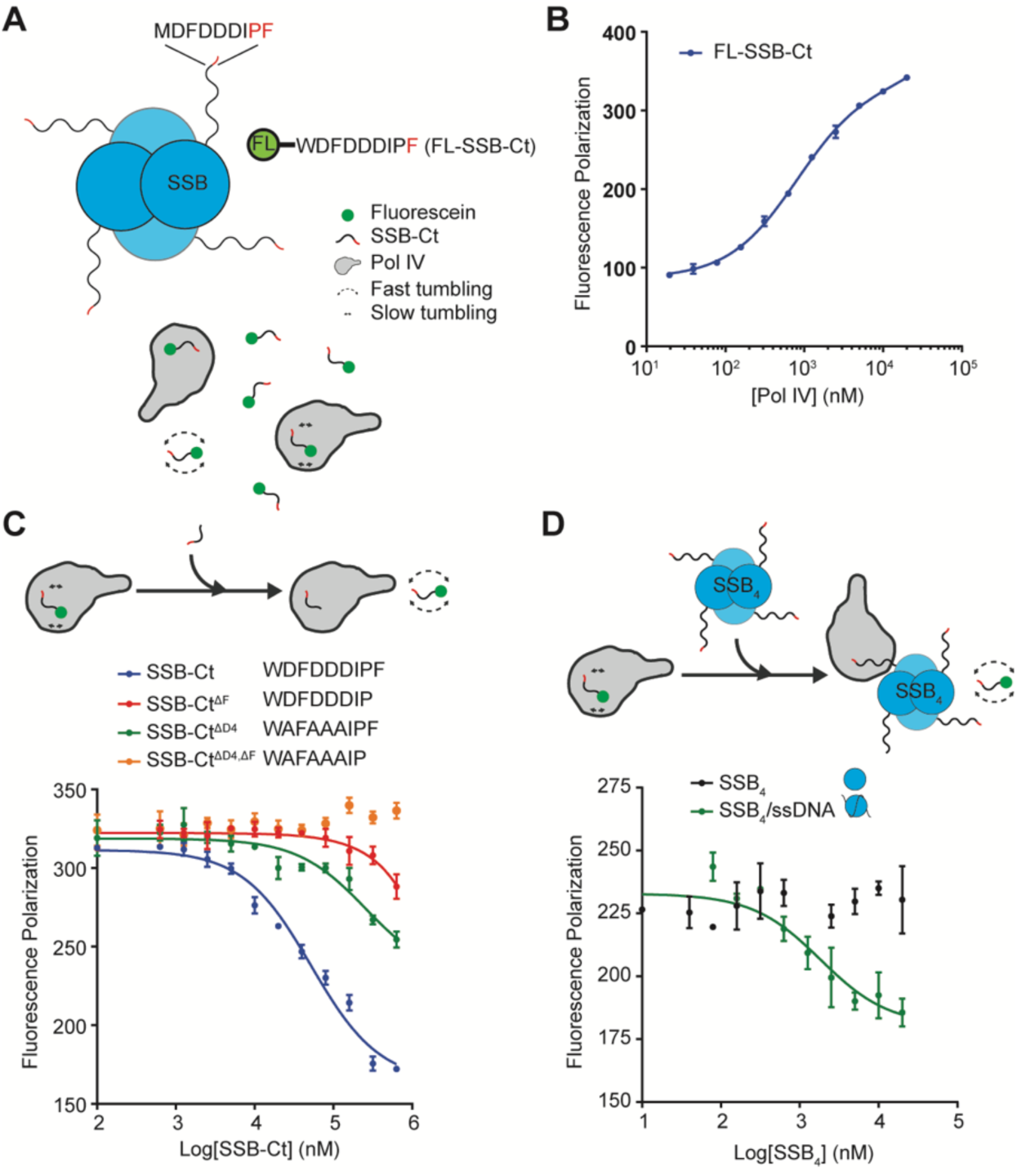
Pol IV selectively interacts with ssDNA-wrapped SSB. A. Fluorescence polarization (FP)-based binding assay. B. Interaction between purified Pol IV and FL-SSB-Ct measured by FP. C. Competitive displacement of FL-SSB-Ct from Pol IV by various unlabeled SSB-Ct peptides. (Top) Competition binding assay scheme. (Bottom) Equilibrium binding isotherms for indicated unlabeled competitors. D. Competitive displacement of FL-SSB-Ct from Pol IV by full-length SSB. (Top) Competition binding assay scheme. (Bottom) Equilibrium binding isotherms.

### Pol IV interacts only with SSB_4_/ssDNA nucleoprotein filaments

To characterize binding of Pol IV to full-length SSB (SSB_4_), we used SSB_4_ as a competitor in the competition binding assays with the FL-SSB-Ct peptide, (Fig 1D, top). Interestingly, in contrast to the isolated SSB-Ct peptide, full-length SSB could not displace bound FL-SSB-Ct from Pol IV (Fig 1D, bottom). The solubility-limited maximum concentration of SSB_4_ (20 μM) used here is equivalent to 80 μM of the isolated SSB-Ct peptide, a concentration, which displaced almost all of the bound FL-SSB-Ct (Fig 1C). These results demonstrate that the binding affinity of bare SSB_4_ to Pol IV is clearly lower than the affinity of the isolated SSB-Ct, and therefore SSB-Ct in SSB_4_ is likely inhibited from interacting with Pol IV. To test whether association of ssDNA with SSB_4_ enabled SSB to interact with Pol IV, we pre-formed SSB_4_/ssDNA complexes by incubating equimolar amounts of SSB_4_ and ssDNA (Fig S2A) and used these complexes as competitor in competition binding assays. Unlike bare SSB_4_, SSB_4_/ssDNA could displace the bound FL-SSB-Ct with Ki of ∼2 μM (Fig 1D), which required the C-terminal ultimate phenylalanine of SSB (Fig S1C), indicating that association of ssDNA with SSB_4_ enabled SSB-Ct to interact with Pol IV. Moreover, the binding affinity is ∼15 fold higher than that of the isolated SSB-Ct. This strengthening may be due to clustering of 4 SSB-Ct peptides within a single SSB_4_, which either increases association and/or decreases dissociation of SSB-Ct. This preferential interaction of Pol IV with SSB_4_/ssDNA suggests that in cells Pol IV interacts primarily with clusters of SSB_4_ associated with ssDNA, for example at the replication fork, as opposed to free SSB_4_ in cytosol.

### The Pol IV/SSB interaction retains unique structural features

The selective interaction of Pol IV with SSB_4_/ssDNA suggests that the SSB-Ct peptides in bare SSB_4_ are mostly in interaction-incompetent states and association of ssDNA with SSB_4_ increases the interaction-competent fraction. We asked if this is a general feature for SIP/SSB interactions. Unlike with Pol IV, bare SSB_4_ could displace a fraction of FL-SSB-Ct bound to Exonuclease I (Exo I) and PriA (Fig S1D), indicating that the SSB-Ct peptide in bare SSB_4_ can interact with both SIPs. Association of ssDNA with SSB_4_ increased the fractional displacement of bound FL-SSB-Ct from both Exo I and PriA with only small increases in the binding affinities (Fig S1D). These results suggest that the SSB-Ct peptides in SSB_4_ are structurally equilibrated between interaction-incompetent and competent states and association of ssDNA with SSB_4_ stabilizes SSB-Ct peptides in more interaction-competent states for these two SIPs without large changes in affinity. Interestingly, the RecQ/SSB interaction was similar to the Pol IV/SSB interaction; 1) relatively low binding affinity of FL-SSB-Ct and 2) selective interaction with SSB_4_/ssDNA (Fig S1B and D). Therefore, it is likely that the SSB-interacting surface(s) within Pol IV, and possibly RecQ, has structural features that do not allow for it to interact with SSB-Ct peptides in interaction-competent states that are populated in the absence of ssDNA.

### N-terminal polymerase domain interacts with SSB

Given that all known SIPs interact with SSB through its SSB-Ct, investigating the physiological role of interaction between a specific SIP with SSB demands identification of mutations within the SIP that disrupt the interaction with SSB(Shereda et al., 2008). To determine which domain of Pol IV bore the SSB-binding surface(s), we used a FP-based assay to measure the affinity between Pol IV and a SSB_4_/ssDNA complex, in which the 3’ end of ssDNA was FAM conjugated (T_71_-FAM) (Fig 2A, top, and S2A, left). The equilibrium binding affinities between Pol IV and the wild-type SSB and SSB^ΔF^-containing complexes (SSB^ΔF^_4_/ssDNA) measured in this assay were nearly identical to affinities determined via the competition-based scheme (Fig 2A, bottom, and S2B). We also observed that the N-terminal polymerase domain (Pol IV^1-230^) bound to SSB_4_/ssDNA with a similar affinity to that of the full-length Pol IV whereas no measurable binding was observed with the C-terminal little finger domain (Pol IV^LF^) (Fig 2A, middle and bottom). These results indicate that the Pol IV^1-230^ contains a binding site(s) for SSB-Ct of SSB.

**Figure 2.**
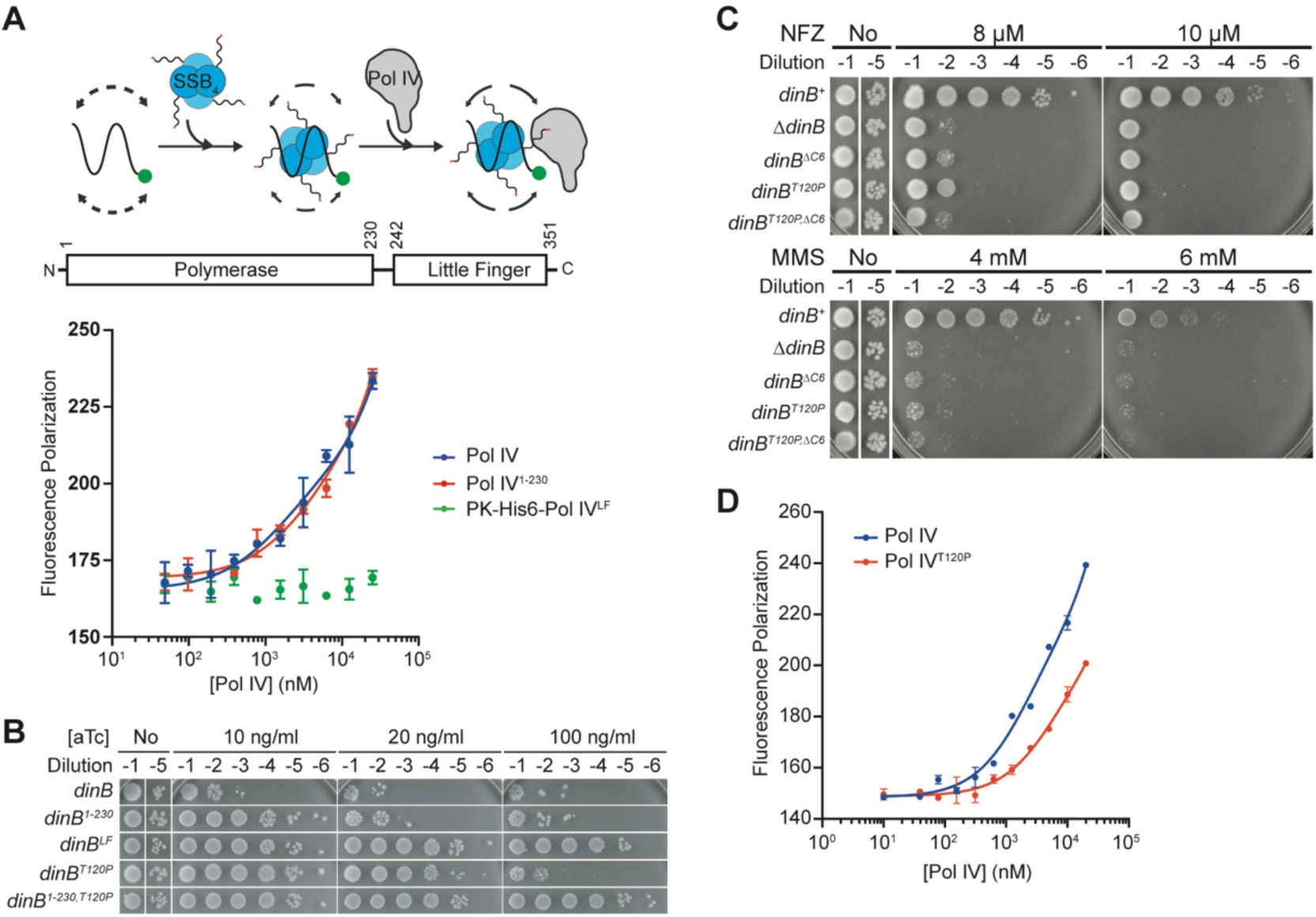
Identification of a SSB-binding-defective mutant Pol IV. A. N-terminal polymerase domain of Pol IV contains a binding site(s) for full-length SSB. (Top) FP-based binding assay using 3’ FAM-labeled ssDNA (T_71_-FAM). (Middle) Domain structure of Pol IV. (Bottom) Interaction of 1) N-terminal polymerase (Pol IV^1-230^) or 2) C-terminal little finger (Pol IV^LF^) domain with SSB_4_/ssDNA. B. The *dinB^T120P^* mutation attenuates cell killing activity of Pol IV. C. Sensitization to NFZ and MMS by the *dinB^T120P^* mutation is epistatic to *dinB^ΔC6^*. D. The *dinB^T120P^* mutation weakens interaction of Pol IV with SSB.

### Leveraging cellular toxicity of over-produced SIPs to discover SSB-binding defective mutations

To discover the SSB-binding surface within the polymerase domain of Pol IV, we exploited the cellular lethality caused by over-expressed Pol IV(Uchida et al., 2008) (Fig 2B). Intriguingly, we observed a similar cell killing effect with other SIPs that we tested (RecQ, Exo I, Topoisomerase III, and Pol II) (Fig S2C). The lethality of over-produced RecQ and Exo I depended on their interaction with SSB as mutations in RecQ (*recQ^R503A^*) (Shereda et al., 2009) and Exo I (*exoI^R184A^*, *exoI^R316A^* and *exoI^Q311A^*) (Lu and Keck, 2008) that reduce binding affinity to SSB attenuated lethality (Fig S2C).

Based on these observations, we hypothesized that the interaction of Pol IV with SSB is required for the lethality by over-produced Pol IV, which would be diminished by Pol IV mutations that weaken the interaction with SSB. Consistent with the polymerase domain of Pol IV containing a binding site(s) for SSB, over-production of the polymerase domain (*dinB^1-230^*) led to massive cell death whereas over-production of the little finger domain (*dinB^LF^*) did not (Fig 2B and S2D). We then surveyed a collection of point mutations within Pol IV that had been previously selected for diminished cell killing activity of Pol IV and found the *dinB^T120P^* mutation within the polymerase domain promising(Scotland et al., 2015) (Fig 2B). Intriguingly, the *dinB^T120P^* mutation influences neither polymerase nor clamp binding activities of Pol IV, yet compromises Pol IV-mediated TLS as the mutation severely sensitized cells to Pol IV cognate damaging agents, which was epistatic to *dinB^ΔC6^* (Fig 2C). These results suggest that Pol IV^T120P^ fails to interact with an unidentified Pol IV-interacting factor, possibly SSB, that regulates the activity of Pol IV in cells.

### The *dinB^T120P^* mutation weakens the interaction between Pol IV and SSB

To examine whether the *dinB^T120P^* mutation indeed weakened the interaction with SSB, we measured the binding of full-length Pol IV^T120P^ to SSB_4_/T_71_-FAM and found that the *dinB^T120P^* mutation reduced the binding affinity ∼3 fold compared to wild-type Pol IV (Fig 2D). Mutating Thr^120^ to proline likely only leads to local structural changes as Pol IV^T120P^ retains wild-type folding and thermal stability (Fig S2E) as well as wild-type polymerase and clamp binding activities(Scotland et al., 2015). Moreover, mutating Thr^120^ to other amino acids also sensitized cells to nitrofurazone (NFZ) to varying degrees (Fig S2F). Among these, mutation to serine (*dinB^T120S^*), whose side chain is structurally closest to Thr, least sensitized cells, while mutation to aspartate (*dinB^T120D^*), which is charged and bulkier than Thr sensitized cells as much as the *dinB^T120P^* mutation did. Collectively, these results suggest that the side chain of Thr^120^ is directly engaged in the interaction with SSB.

### SSB facilitates access of Pol IV to the replication fork

Given Pol IV^T120P^ was severely compromised in mediating TLS in cells, we asked whether the Pol IV-SSB interaction might enable Pol IV to access the replication fork. To address this question, we exploited the replication-inhibitory action of Pol IV on the *in vitro* reconstituted *E. coli* replisome, which results from the relatively slow Pol IV substituting for Pol III in replication(Indiani et al., 2009; Uchida et al., 2008) (Fig 3A).

**Figure 3.**
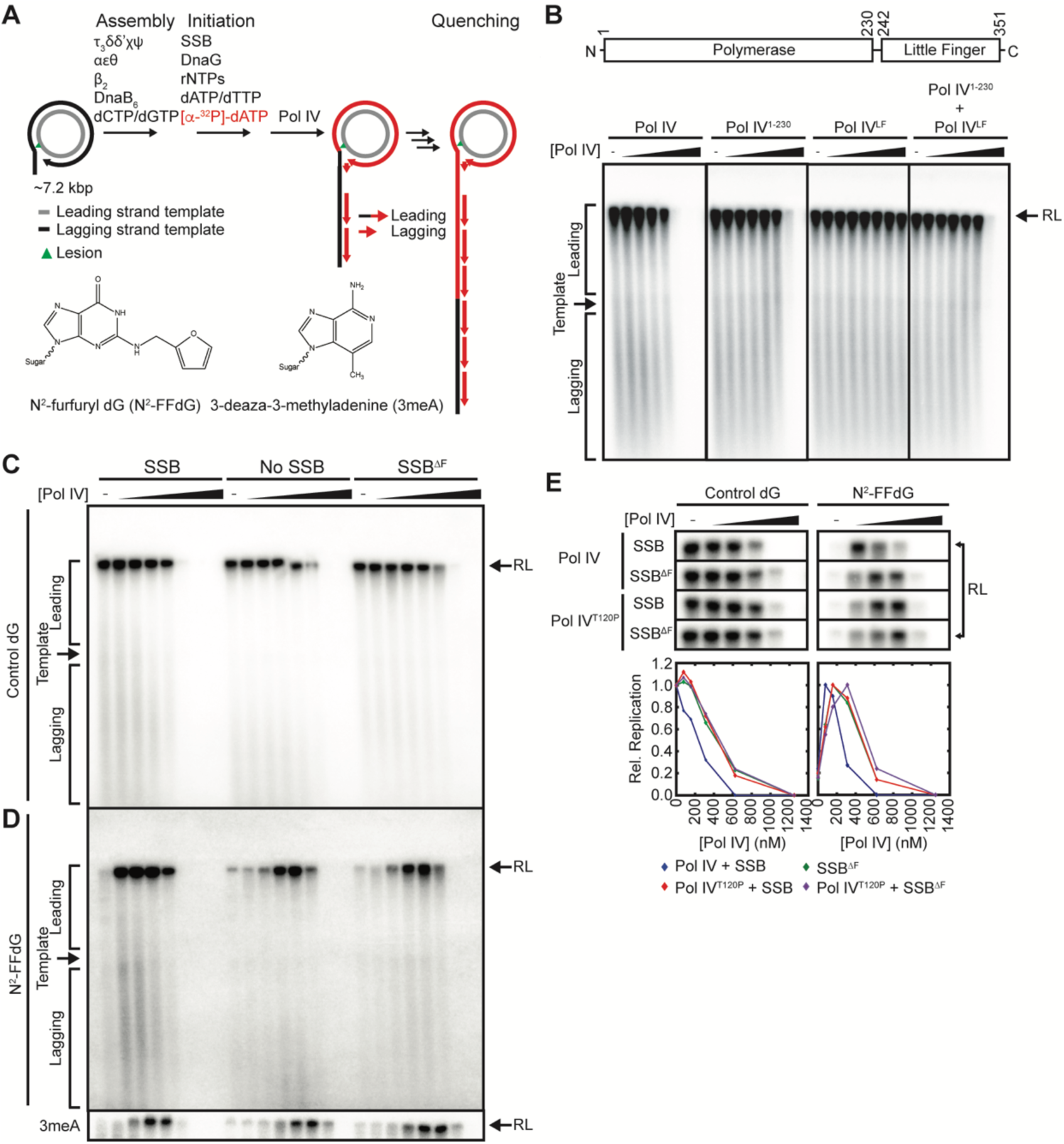
SSB facilitate access of Pol IV to the replication fork. A. *In vitro* reconstitution of Pol IV-mediated TLS on a rolling circle template that bears either N^2^-FFdG or 3meA on the leading strand template. Replication products were separated on an alkaline agarose gel (0.6%) and visualized by autoradiography. Long leading strand replication products accumulate as a band at the resolution limit (RL) of the gel (∼45 kilonucleotides) while short lagging strand replication products run as diffusive bands below the unreplicated template (see panel B). B. N-terminal polymerase domain of Pol IV is sufficient for replication-inhibitory activity of Pol IV. (Top) Domain structure of Pol IV. (Bottom) A lesion-free control template was replicated in the presence of indicated Pol IV variants (39/78/156/312/625/1250/2500 nM final), which were added 10 sec after the initiation of replication. C. Interaction of SSB with Pol IV potentiates replication-inhibitory activity of Pol IV. A lesion-free control template was replicated either in the presence of SSB or SSB^ΔF^, or in the absence of SSB. Indicated amounts of Pol IV were added 10 sec after the initiation of replication. D. Interaction of SSB with Pol IV potentiates Pol IV-mediated TLS over N^2^-FFdG or 3meA. The N^2^-FFdG- or 3meA-containing template was replicated either in the presence of SSB or SSB^ΔF^, or in the absence of SSB. Varying amounts of Pol IV (39/78/156/312/625/1250/2500 nM final) were added 10 sec after the initiation of replication. Refer to Fig S3A for the entire gel showing replication products of a 3meA-containing template. E. The *dinB^T120P^* and *ssb^ΔF^* mutations redundantly attenuate both replication-inhibitory and TLS activities of Pol IV. A control template (left) or a N^2^-FFdG-containing template was replicated in the presence of SSB or SSB^ΔF^. Varying amounts (78/156/312/625/1250 nM final) of Pol IV or Pol IV^T120P^ were added 10 sec after the initiation of replication.

Consistent with prior observations(Chang et al., 2019), addition of increasing amounts of full-length Pol IV to replication reactions on a rolling-circle template led to gradual reductions in both leading and lagging strand synthesis (Fig 3B). Previous studies largely attributed this inhibitory action to the little finger domain (Pol IV^LF^) because it bears clamp-binding activity and thus could displace Pol III from the β_2_ clamp. However, we observed that isolated Pol IV^LF^ could not inhibit replication (Fig 3B). Rather, the isolated polymerase domain (Pol IV^1-230^) could inhibit replication but less potently compared with the full-length Pol IV. Moreover, addition of both Pol IV^1-230^ and Pol IV^LF^ resulted in a similar reduction of replication to that by Pol IV^1-230^ alone. These results suggest a cooperative action in inhibiting replication of Pol IV^1-230^ and Pol IV^LF^ within full-length Pol IV. Given Pol IV^1-230^ has SSB-binding activity (Fig 2A), it is possible that Pol IV^1-230^ potentiates clamp-binding activity of Pol IV by interacting with SSB.

If SSB promotes access of Pol IV to the replication fork, omitting SSB in replication reactions should attenuate the replication-inhibitory action of Pol IV. Indeed, in the absence of SSB, Pol IV inhibited replication 2 to 3 fold less potently as compared with inhibition in the presence of SSB (Fig 3C). Replacing SSB with SSB^ΔF^, which is selectively defective in interacting with Pol IV but maintains wild-type ssDNA binding, similarly attenuated the replication-inhibitory action of Pol IV (Fig 3C), suggesting that SSB directly potentiates the action of Pol IV at the fork. Notably, given omitting SSB and replacing SSB with SSB^ΔF^ resulted in similar effects on Pol IV-inhibition of replication, it is likely that the transient Pol IV-SSB interaction potentiates action of Pol IV by locally concentrating free Pol IV molecules near replisomes rather than the formation of a more potent stable Pol IV-SSB binary complex.

### SSB facilitates access of Pol IV to the lesion-stalled replisomes

Next, we asked if the Pol IV-SSB interaction also facilitates access of Pol IV to lesion-stalled replisomes. For this we introduced a single N^2^-FFdG into the leading strand template (Fig 3A), which potently blocked synthesis of both leading and lagging strands(Chang et al., 2019) (No Pol IV in Fig 3D). Upon addition of increasing concentrations of Pol IV to replication reactions of the lesion-containing template, synthesis of both leading and lagging strands was gradually restored before Pol IV completely inhibited replication at high concentrations (Fig 3D). The robust accumulation of resolution-limited (RL) leading strand replication products indicates that Pol IV mediates TLS over the N^2^-FFdG adduct very efficiently without creating strand discontinuities, such as ssDNA gaps and nicks that would terminate replication upon subsequent passage around the template(Chang et al., 2019) (Fig S3A). When SSB was omitted, Pol IV similarly restored both leading and lagging strand synthesis but 2 to 3 fold higher concentrations of Pol IV were required for peak TLS as compared with peak TLS in the presence of SSB (Fig 3D). The apparent reduction in the amount of leading and lagging strand replication products in the absence of SSB was likely attributable to the general reduction in processive replication (Fig 3C). However, replication-normalized peak TLS in the absence of SSB was comparable to that in the presence of SSB, indicating that the Pol IV-SSB interaction does not make Pol IV more efficient in mediating TLS over a lesion. We made similar observations with a 3meA-containing template (Fig 3A and D and S3B), but it is noteworthy that Pol IV is less efficient in replicating past 3meA than N^2^-FFdG as higher concentrations of Pol IV were required for TLS over 3me-dA compared with N^2^-FFdG. Replacing SSB with SSB^ΔF^ resulted in similar observations (Fig 3D and S3B), indicating that SSB in lesion-stalled replisomes potentiates Pol IV-mediated TLS by facilitating the access of Pol IV to the replication fork rather than making Pol IV more efficient in mediating TLS by forming a stable Pol IV-SSB binary complex.

### The *dinB^T120P^* mutation reduces access of Pol IV to advancing or stalled replisomes

We next tested whether the *dinB^T120P^* mutation had similar effects on processive replication and TLS to those of SSB^ΔF^, which would indicate that compromised TLS in the *dinB^T120P^* strain was due to the defective interaction of Pol IV^T120P^ with SSB. Similar to wild-type Pol IV, Pol IV^T120P^ inhibited processive replication but 2 to 4 fold less potently, which resembled the effect of replacing SSB with SSB^ΔF^ (Fig 3E, left). Importantly, replacing SSB with SSB^ΔF^ did not further reduce the potency of Pol IV^T120P^ (Fig 3E, left), indicating that the *dinB^T120P^* and *ssb^ΔF^* mutations acted redundantly. Similarly, Pol IV^T120P^ mediated TLS over the N^2^-FFdG adduct with 2 to 4 fold reduced potency as compared with wild-type Pol IV (Fig 3E, right), yet Pol IV^T120P^ retained nearly wild-type efficiency in mediating TLS. Similar to the effect on the replication-inhibitory action of Pol IV, replacing SSB with SSB^ΔF^ barely reduced the potency of Pol IV^T120P^ (Fig 3E, right). Collectively these results demonstrate that reduced access of Pol IV^T120P^ to the replication fork for both advancing and lesion-stalled replisomes is due to its reduced binding to SSB.

### The *dinB^T120P^* mutation abolishes damage-induced enrichment of Pol IV near replisomes in living cells

Given that the *dinB^T120P^* mutation compromises access of Pol IV to the replication fork *in vitro*, we speculated that the interaction between SSB and Pol IV plays a critical role in localizing Pol IV to stalled replisomes in cells. To explore this possibility, we employed single-particle tracking PALM (Photoactivated Localization Microscopy) imaging in living cells to track the location of individual Pol IV molecules. We replaced the endogenous copy of Pol IV with a fusion to PAmCherry, a photoactivatable fluorescent protein (Pol IV-PAmCherry) (Thrall et al., 2017) (Fig 4A and B) and carried out imaging in strains containing the *lexA(Def)* mutation, which constitutively derepresses SOS-response genes. As PAmCherry is initially in a non-fluorescent dark state, we used a low level of continuous near-UV excitation to stochastically activate Pol IV-PAmCherry molecules, which could then be imaged with excitation at 561 nm until they photobleached. The UV excitation power was optimized to ensure that on average only a single Pol IV-PAmCherry molecule was activated at any one time in a given cell. In prior work we showed that there are two populations of Pol IV, a fast diffusing population and a static population(Thrall et al., 2017). The static population represents Pol IV molecules that are bound to DNA, either associated with the replisome or elsewhere in the cell. In order to assess association of Pol IV with replisomes, we first determined the locations of replisomes using SSB-mYPet(Thrall et al., 2017). Subsequently, in the same cells, PALM imaging was carried out in a way that selectively resolved static Pol IV-PAmCherry populations (Fig. 4B). We then calculated the distance of each static Pol IV molecule to the nearest SSB focus. Consistent with our prior observations(Thrall et al., 2017), upon treatment of cells with MMS (100 mM), the Pol IV-SSB distance distribution shifted to shorter distances (Fig 4C), indicative of more Pol IV molecules colocalizing with replisomes. To quantify enrichment of Pol IV near the replisome, the distance distribution between Pol IV and SSB foci was further analyzed with a radial distribution function, g*(r)*, which expresses the likelihood of Pol IV being found within a distance *r* from SSB relative to random cellular localization; values of g*(r)* greater than 1 indicate enrichment (Fig S4A). In MMS-treated cells, Pol IV was about 8 fold enriched (g*(r)* ∼ 8) near replisomes relative to random localization (Fig 4D, and S4B and C), whereas it was barely enriched in untreated cells(Thrall et al., 2017). Notably, unlike wild-type Pol IV, upon treatment with MMS, the distance distribution between Pol IV^T120P^ and SSB foci barely shifted to shorter distances (Fig 4C), and Pol IV^T120P^ was only about 2 fold enriched near replisomes (Fig 4D). These results indicate that the Pol IV-SSB interaction plays a dominant role in the localization of Pol IV to stalled replisomes. The residual enrichment (g(*r*) ∼ 2) may be attributable to the residual SSB-binding activity of Pol IV^T120P^ (Fig 2D). This notion is also consistent with the *dinB^T120P^* mutation being hypomorphic; Pol IV-mediated TLS in the *dinB^T120P^* strain is severely attenuated but less defective than that in the *ΔdinB* strain (Fig. 2C). Considering purified Pol IV^T120P^ retained a nearly wild-type thermal stability and folding (Fig. S2E), comparable expression between Pol IV and Pol IV^T120P^ in the imaging strains (Fig S4D) rule out abnormal behaviors of Pol IV^T120P^ in cells, such as aggregation.

**Figure 4.**
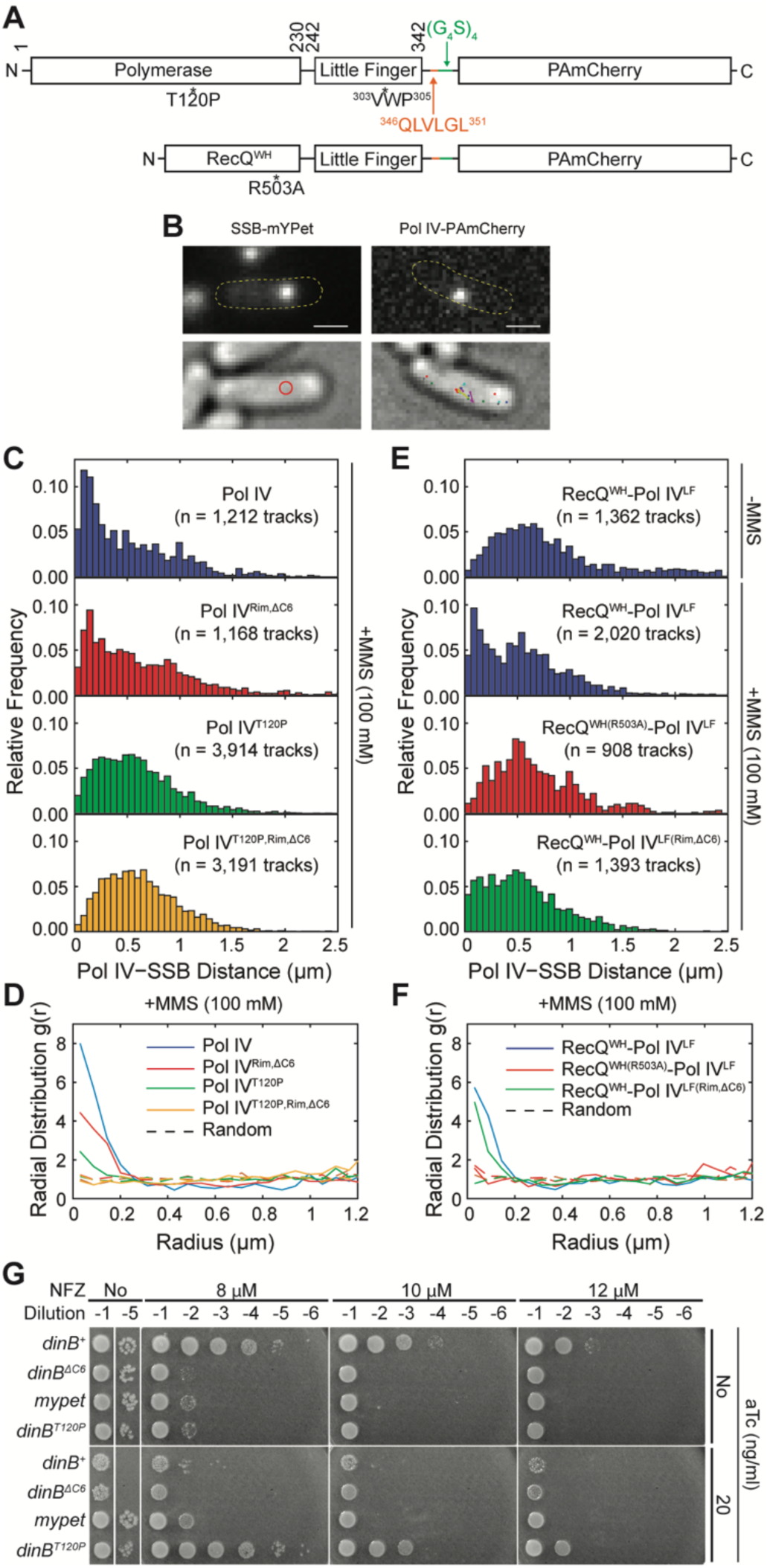
The Pol IV-SSB interaction enriches Pol IV near lesion-stalled replisomes in cells. A. Schematic diagrams of PAmCherry fusions of Pol IV and RecQ^WH^-Pol IV^LF^ variants. ^303^VWP^305^, rim interacting residues; ^346^QLVLGL^351^, the CBM of Pol IV; (G_4_S)_4_, a flexible linker. B. (Top panels) Representative fluorescence micrographs of an SSB-mYpet focus (left) and a single photoactivated Pol IV-PAmCherry molecule (right) with overlays showing the cell outlines. (Bottom panels) Corresponding brightfield micrographs with overlays showing SSB foci (red circle, left) or all detected Pol IV tracks (right) (Scale bars: 1 µM). C. Distributions of the mean distance between each static Pol IV track and the nearest SSB focus for Pol IV^WT^ and mutants in cells treated with 100 mM MMS. D. Radial distribution functions *g(r)* for Pol IV^WT^ and mutants in cells treated with 100 mM MMS. Also shown are random *g(r)* functions for each data set (dotted lines). E. Distributions of the mean distance between each static RecQ^WH^-Pol IV^LF^ track and the nearest SSB focus for wild-type RecQ^WH^-Pol IV^LF^ and mutants in cells either untreated or treated with 100 mM MMS. F. Radial distribution functions *g(r)* RecQ^WH^-Pol IV^LF^ and mutants in cells treated with 100 mM MMS. Also shown are random *g(r)* functions for each data set (dotted lines). G. Over-production of Pol IV^T120P^ restores wild-type tolerance to NFZ in a *ΔdinB* strain. (Top) A tet-inducible expression cassette engineered within the *lamB* gene in a *ΔdinB* strain. (Bottom) Serially diluted cultures of strains with indicated genes in the tet-inducible expression cassette were spotted on LB-agar plates containing varying concentrations of NFZ and anhydrous tetracycline (aTc). Over-production of neither Pol IV^DC6^ nor mYpet restores tolerance to NFZ. Refer to Fig S4E for lower aTc concentrations.

Intriguingly mutating both the CBM and rim residues (Pol IV *^Rim,Δ^*^C6^), which completely abolishes interaction of Pol IV with the β_2_ clamp and thus Pol IV-mediated TLS(Chang et al., 2019), only partially reduced the MMS-induced enrichment of Pol IV near replisomes(Thrall et al., 2017) (Figure 4C and 4D). Combining the *dinB^T120P^* and *dinB^Rim,ΔC6^* mutations completely abolished enrichment of Pol IV near replisomes (Figure 4C and 4D), indicating that the MMS-induced enrichment of Pol IV is fully attributable to the interactions of Pol IV with SSB and the β_2_ clamp. However, the combined change in magnitude in g*(r)* (∼10) upon individually ablating the Pol IV-SSB (Pol IV^T120P^) and Pol IV-β_2_ clamp (Pol IV^Rim,ΔC6^) interactions is greater than the value of g*(r)* for wild-type Pol IV (∼7) indicating that these interactions do not independently contribute to the enrichment of Pol IV near replisomes. Given the effect of the *dinB^T120P^* mutation on colocalization of Pol IV with the replisome is substantially greater than the effect of the *dinB^Rim,ΔC6^* mutation, it is possible that the Pol IV-SSB interaction plays a dominant role in localizing/enriching Pol IV near stalled replisomes and thus poises Pol IV to efficiently associate with the β_2_ clamp, allowing for TLS. In this model, the partial reduction in the enrichment of Pol IV by the *dinB^Rim,ΔC6^* mutation could reflect the disappearance of the β_2_ clamp-bound fraction of Pol IV that performs TLS rather than reduced localization of Pol IV to replisomes.

### Interaction of Pol IV with SSB is sufficient for damage-induced enrichment of Pol IV near replisomes

Our observation suggests that interaction of Pol IV with SSB is sufficient to localize Pol IV near replisomes in MMS-treated cells. However, Pol IV is known to interact with other factors(Cafarelli et al., 2014; Cohen et al., 2009; Godoy et al., 2007; Sladewski et al., 2011) including UmuD and RecA and these factors may play a role in localizing Pol IV to replisomes. To examine if the interaction with SSB is sufficient for damage-induced enrichment of Pol IV near replisomes, we replaced the polymerase domain of Pol IV (Pol IV^1-230^) with the SSB-interacting winged helix (WH) domain of the *E. coli* RecQ (RecQ^WH^), creating a chimeric protein, RecQ^WH^-Pol IV^LF^ (Fig 4A). We chose RecQ^WH^ because 1) RecQ^WH^ bears a well characterized binding site for SSB-Ct with no known enzymatic activities and 2) RecQ^WH^ is not known to interact with Pol IV-interacting factors. Given that the polymerase domain of Pol IV bears a SSB-binding site(s) and interacts with all the known Pol IV-interacting proteins but the β_2_ clamp, damage-induced enrichment of this chimeric protein would demonstrate that interaction with SSB is sufficient to localize Pol IV to stalled replisomes. This chimeric protein was expressed from the native *dinB* locus as a C-terminal fusion to PAmCherry (RecQ^WH^-Pol IV^LF^-PAmCherry), resulting in the imaging strain lacking Pol IV. Indeed, similar to the wild-type Pol IV, the chimeric protein was highly enriched (*g(r)* ∼ 6) near replisomes in MMS-treated cells but not in untreated cells (Fig 4E and 4F). Importantly, similar to the effect of the *dinB^T120P^* mutation on the localization of Pol IV, a mutation within the RecQ^WH^ domain (RecQ^WH(R503A)^-Pol IV^LF^), which reduces the binding affinity of RecQ^WH^ to the SSB(Shereda et al., 2009), nearly completely abolished the damage-induced enrichment (Fig 4E and 4F). These results clearly demonstrate that interaction of Pol IV with SSB is sufficient for localizing Pol IV to lesion-stalled replisomes and interaction with other factors including the β_2_ clamp does not play a necessary role in this process. However, we do not rule out the possibility that association of Pol IV with the β_2_ clamp, following the SSB-dependent localization to the replisome, could be modulated by other factors, such as UmuD and RecA(Cafarelli et al., 2014).

Notably, unlike wild-type Pol IV, mutating β_2_ clamp-interacting residues in the Pol IV^LF^ domain of RecQ^WH^-Pol IV^LF^ (RecQ^WH^-Pol IV^LF(Rim,ΔC6)^) did not lead to a discernable reduction in the enrichment of the chimeric protein in MMS-treated cells (Fig 4E and 4F). Interaction of a DNA polymerase with a processivity factor or the P/T junction cooperatively enhances both interactions. Unlike Pol IV, interaction between RecQ^WH^-Pol IV^LF^ and β_2_ clamp cannot be enhanced by RecQ^WH^ because RecQ^WH^ is incapable of capturing the P/T junction. This can substantially reduce the lifetime of β_2_ clamp-bound RecQ^WH^-Pol IV^LF^, lowering the chance that this population of RecQ^WH^-Pol IV^LF^ is captured in our imaging scheme. Consistent with this notion, the *g(r)* value for RecQ^WH^-Pol IV^LF^ (∼6) was close to that for Pol IV^Rim,ΔC6^ (∼5) in MMS-treated cells (Fig 4D and F). This observation again supports the notion that interaction of Pol IV with β_2_ clamp does not play a major role in localizing Pol IV to the replisome.

### Elevation of the expression level of Pol IV^T120P^ restores wild-type TLS in cells

Loss of MMS-induced enrichment of Pol IV near replisomes in the *dinB^T120P^* background raises the possibility that SSB enables Pol IV-mediated TLS by increasing local concentrations of Pol IV near lesion-stalled replisomes. In this case, simply elevating the level of Pol IV^T120P^ would restore tolerance to NFZ. To examine this possibility, a tetracycline-inducible expression cassette, from which transcription of either *dinB^+^* or *dinB^T120P^* could be induced by anhydrotetracycline (aTc), was engineered into the genome of a *ΔdinB* strain (Fig S4E, refer to “materials and methods” for details). In the absence of aTc, the strain bearing the wild-type *dinB* (*dinB^+^*) in the expression cassette retained nearly wild-type tolerance to NFZ without exhibiting growth defect whereas the strain bearing *dinB^ΔC6^* or *mype*t in the cassette was highly sensitized to NFZ (Fig 4G and S4E). This inducer-independent complementation of the *ΔdinB* strain indicates a low level of leaky expression of Pol IV. Under the same condition, the strain bearing *dinB^T120P^* (*dinB^T120P^*) was severely sensitized to NFZ but displayed no growth defects, recapitulating the growth and sensitivity of the *dinB^T120P^* strain (Fig 4G). These observations demonstrate that the inducible expression system can reconstitute Pol IV-mediated TLS in the *ΔdinB* strain.

As anticipated, over-production of wild-type Pol IV in the presence of aTc (>10 ng/ml) led to massive cell death (No NFZ in Fig 4G) and also led to reduced tolerance to NFZ. However, this apparent reduction in tolerance likely reflects cell death caused by over-produced levels of Pol IV rather than indicating that Pol IV-mediated TLS per se became compromised. In stark contrast to Pol IV, as the concentration of aTc increased, over-produced Pol IV^T120P^ gradually restored tolerance to NFZ without inhibiting cell growth, which was comparable to the tolerance of the *dinB^+^* strain at optimal concentrations of aTc (Fig 4G and S4E). Collectively, these results demonstrate that the local concentration of Pol IV near lesion-stalled replisomes in *dinB^+^* strains is higher than the average cellular concentration of Pol IV due to the SSB-mediated enrichment.

### Pol IV-SSB interaction facilitates polymerase switching within lesion-stalled replisomes

As TLS at the fork competes with repriming of DNA synthesis downstream of the lesion(Chang et al., 2019), we asked whether the defect in TLS of Pol IV^T120P^ leads to an increase in repriming. Similar to Okazaki fragments, repriming products are shorter than the template in our rolling circle replication assay (Fig S5A). However, unlike Okazaki fragments, repriming products result from leading strand synthesis and can be selectively probed by Southern blotting with leading strand specific probes(Chang et al., 2019) (Fig 5A, top). In the absence of Pol IV, a fraction of N^2^-FFdG-stalled replisomes reprimed DNA synthesis, forming short leading strand replication products (Fig 5A, bottom). Long leading strand replication products, which can only be produced when TLS at the fork happens over the N^2^-FFdG lesion, were also formed due to inefficient replication of Pol III past N^2^-FFdG(Chang et al., 2019) (Fig 5A). Addition of increasing amounts of wild-type Pol IV to replication reactions led to a gradual increase in resolution limited products, which result from TLS at the fork (Figure 5A, bottom). This increase in TLS was accompanied by a concomitant decrease in repriming, consistent with a kinetic competition between both pathways(Chang et al., 2019). Compared with wild-type Pol IV, Pol IV^T120P^ mediated TLS 2 to 4 fold less potently, and repriming persisted to 2 to 4 fold higher concentrations (Fig 5A, bottom). We also made similar observations with the 3meA-containing template although repriming persisted to higher concentrations of Pol IV, presumably because Pol IV-mediated TLS over 3meA is less efficient as compared to N^2^-FFdG (Fig 5A, bottom). This persistent repriming reflects inefficient polymerase switching from Pol III to Pol IV^T120P^ within lesion-stalled replisomes.

**Figure 5.**
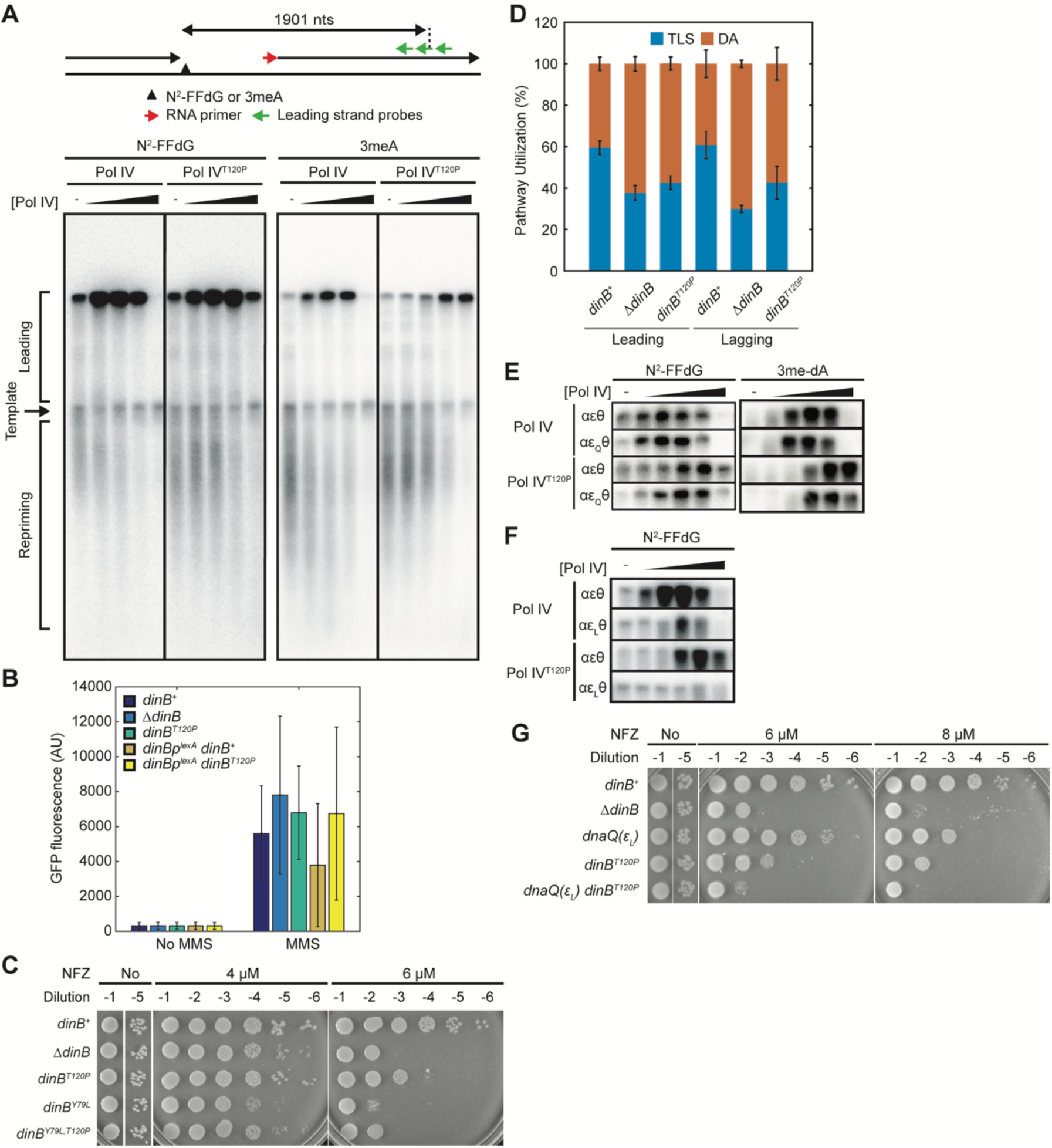
Local enrichment of Pol IV near stalled replisomes promote TLS at the fork. A. The SSB-Pol IV interaction suppresses repriming by promoting TLS. (Top) Repriming of DNA replication by a lesion-stalled replisome, and Southern blot probes for detecting repriming products. (Bottom) Attenuated TLS by the *dinB^T120P^* mutation results in persistent repriming. Replication products were detected by Southern blotting with a cocktail of radiolabeled leading strand probes. B. The MMS-induced SOS response is elevated in the *dinB^T120P^* strain. SOS-response reporter strains bearing indicated *dinB* alleles were treated with 6 mM MMS for 1 hr and GFP fluorescence from single cells was measured by flow cytometry. Results are averages and standard deviations of 10^5^ cells. C. Hypersensitivity of a *dinB^Y79L^* strain to NFZ is alleviated by the *dinB^T120P^* mutation. D. The *dinB^T120P^* mutation compromises TLS over the N^2^-FFdG adduct on both leading and lagging strand templates in cells. E. Weakening the ε-β_2_ clamp interaction potentiates Pol IV^T120P^-mediated TLS. F. Strengthening the ε-β_2_ clamp interaction prevents Pol IV^T120P^ from mediating TLS. G. Synthetic suppression of TLS by the *dinB^T120P^* and the *dnaQ(ε_L_)* mutations. Sensitivity to NFZ of indicated strains.

### SSB facilitates polymerase switching within stalled replisomes in cells

We next examined if repriming was favored over TLS at the fork in cells bearing the *dinB^T120P^* mutation. Repriming results in the formation of ssDNA gaps that activate the SOS DNA damage response(Chang et al., 2019; Yeeles and Marians, 2013) (Fig S5A). To measure induction of this response, we created SOS-response reporter strains, in which expression of GFP is controlled by the promoter of the *sulA* gene, a tightly repressed SOS responsive gene(Chang et al., 2019; McCool et al., 2004). Upon MMS treatment, expression of GFP in the reporter strain with the wild-type *dinB* was highly induced (Fig 5B), indicating the creation of ssDNA gaps. Consistent with the repriming-suppressive effect of Pol IV *in vitro*, constitutive de-repression of *dinB* by mutating the LexA repressor binding site in the *dinB* promoter (*dinBp^lexA^ dinB^+^*) reduced the MMS-induced SOS response, whereas deleting the *dinB* gene (*ΔdinB*) elevated the MMS-induced SOS response(Chang et al., 2019) (Fig 5B). Notably, the MMS-induced SOS response was elevated in the *dinB^T120P^* strain compared with the wild-type *dinB* strain (*dinB^+^*). Moreover, in contrast to wild-type *dinB*, de-repression of *dinB^T120P^* did not suppress the MMS-induced SOS response (Fig 5B). Collectively, these results suggest that a higher fraction of lesion-stalled replisomes reprime DNA replication in the *dinB^T120P^* strain than in the *dinB^+^* strain.

More frequent repriming in the *dinB^T120P^* strain suggests inefficient polymerase switching from Pol III to Pol IV within lesion-stalled replisomes. To explore this possibility, we exploited the *dinB^Y79^* allele, which sensitizes cells to damaging agents even more so than the *dinB* deletion allele(Benson et al., 2014; Jarosz et al., 2009). This hypersensitization results from Pol IV^Y79L^ acting as a suicide inhibitor of the stalled replisome; Pol IV^Y79L^ switches normally with Pol III but cannot complete TLS, thereby suppressing both Pol IV- and Pol III-mediated TLS(Chang et al., 2019). If the *dinB^T120P^* mutation prevents Pol IV from switching with Pol III, the *dinB^T120P^* mutation should mitigate the hypersensitivity of the *dinB^Y79L^* strain to damaging agents. Indeed, the *dinB^T120P^* mutation lessened the hypersensitivity of *dinB^Y79L^* (*dinB^Y79L,T120P^*) to NFZ (Fig. 5C). Collectively, these results indicate that in the *dinB^T120P^* strain, polymerase switching within lesion-stalled replisomes occurs inefficiently, diverting more lesion-stalled replisomes into the repriming pathway.

### In vivo TLS on both leading and lagging strands is compromised by the *dinB^T120P^* mutation

To directly evaluate the effect of the *dinB^T120P^* mutation on Pol IV-mediated TLS in cells, we employed an *in vivo* TLS assay that quantitatively measures fractions of lesion-stalled replisomes resolved through either TLS or homology-dependent damage avoidance (DA) pathways in a strand specific manner(Pagès et al., 2012). In this assay, replication blocking DNA lesions are site specifically introduced into the *E. coli* genome (Fig S5B, left). Here, we introduced into either the leading-strand or lagging-strand templates of the *E. coli* genome, a single N^2^-furfuryl dG (N^2^-FFdG) lesion, which is a structural analogue of DNA lesions created in NFZ-treated *E. coli* cells(Jarosz et al., 2009; 2006). In *E. coli*, these lesions are efficiently removed by nucleotide excision repair (NER) (Ona et al., 2009), and in order to keep the lesion, we used NER-deficient strains, in which *uvrA* gene was knocked out. The *E. coli* strains used in the assay were engineered to express functional LacZ only when replisomes stalled at replication-blocking lesions are released by TLS, resulting in formation of blue-sectored colonies on X-gal containing plates (Fig S5B, right).

Replisomes stalled at N^2^-FFdG on either the leading-strand or lagging-strand templates were resolved by a combination of TLS and DA (Fig 5D). Among cells that tolerated the N^2^-FFdG lesion and formed colonies, about 60% for both the leading and the lagging strand lesions were blue-sectored, indicating that the majority of stalled replisomes at N^2^-FFdG were resolved by TLS (Fig 5D). Consistent with the genetic requirement of the *dinB* gene for the tolerance to NFZ(Jarosz et al., 2009) (Fig 2C), deletion of the *dinB* gene (*ΔdinB*) reduced the fraction of stalled replisomes on both the leading and lagging strand templates that were resolved by TLS to about 30% with concomitant increases in the utilization of the DA pathway (Fig 5D). The residual TLS in the *ΔdinB* background (Fig S5C and D) was likely due to inefficient but measurable Pol III-mediated TLS over N^2^-FFdG(Chang et al., 2019).

In the *dinB^T120P^* background, about 40% of stalled replisomes on the leading strand template were resolved by TLS, which is only slightly higher than in *ΔdinB*, demonstrating that the *dinB^T120P^* mutation severely compromises Pol IV-mediated TLS in cells. This severe but partial defect is consistent with the *dinB^T120P^* mutation weakening the interaction of Pol IV with SSB but not completely ablating the interaction (Fig 2D). Intriguingly, utilization of TLS at N^2^-FFdG on the lagging strand template was similarly reduced by the *dinB^T120P^* mutation. Similar utilization of TLS and impact of the *dinB^T120P^* mutation in resolving stalled replisomes on the leading and lagging strand templates suggests that a common regulatory mechanism controls pathway utilization on both strands.

### SSB-dependent enrichment enables Pol IV to overcome the ε kinetic barrier

We previously showed that the frequency of repriming increased, at the expense of TLS at the fork, when the interaction between the ε subunit of Pol III core and the β_2_ clamp was strengthened(Chang et al., 2019). This interaction acts as a molecular gate to regulate clamp binding and thus strengthening it suppresses association of Pol IV with the β_2_ clamp, a prerequisite for Pol IV mediating TLS. Our observation that Pol IV enrichment near stalled replisomes is required for Pol IV-mediated TLS suggests that a relatively high local concentration of Pol IV is necessary for Pol IV to compete with the ε subunit for clamp binding. If this is the case, weakening the ε-cleft interaction, which reduces the TLS-inhibitory activity of the ε subunit(Chang et al., 2019), would potentiate the action of Pol IV^T120P^. Indeed, when the ε-β_2_ clamp interaction was weakened by the *dnaQ(ε_Q_)* mutation (αε_Q_θ), which reduces binding affinity of the ε subunit to the β_2_ clamp by >500 fold(Jergic et al., 2013) (Fig S5E), Pol IV^T120P^ could mediate TLS over both N^2^-FFdG and 3meA at lower concentrations than it did within the wild-type replisome (αεθ) (Fig 5E).

Conversely, when the ε-cleft interaction was strengthened by the *dnaQ(ε_L_)* mutation, which modestly suppresses wild-type Pol IV-mediated TLS(Chang et al., 2019) (Fig S5E), Pol IV^T120P^ barely mediated TLS over N^2^-FFdG (Fig 5F). Consistent with the *in vitro* synthetic suppression, the strain bearing both the *dinB^T120P^* and the *dnaQ(ε_L_)* mutations [*dnaQ(ε_L_) dinB^T120P^*] was as severely sensitized to NFZ as the *dinB* deletion strain (*ΔdinB*), while the strain bearing either the *dinB^T120P^* or the *dnaQ(ε_L_)* mutation was only modestly sensitized(Chang et al., 2019) (Fig 5G). Collectively, these results demonstrate that in cells Pol IV must be highly enriched near stalled replisomes to overcome the kinetic barrier imposed by the ε subunit(Chang et al., 2019).

### CLIPs devoid of SSB binding activity are excluded from the replication fork

Given that SSB-mediated enrichment of Pol IV is required to overcome the ε kinetic barrier, we asked whether other CLIPs require SSB binding to access the β_2_ clamp at the replication fork. Unlike Pol IV, over-production of Pol I and Crfc, CLIPs that lack SSB-binding activity, did not cause cell death (Fig 6A). However, over-production of Pol I (*polA*) fused to the SSB binding domain of RecQ (Pol I-RecQ^WH^) caused massive cell death (Fig 6A) whereas Pol I-RecQ^WH(R425A,R503A)^, which lacks affinity for SSB, failed to kill cells. We also made similar observations with Crfc, a CLIP that lacks polymerase activity (Fig 6A). Cell killing by Pol I-RecQ^WH^ or Crfc-Rec^WH^ required both SSB and clamp binding activities because overproduction of RecQ^WH^ alone did not kill cells (Fig S2F). Similarly, overproduction of Pol IV^LF^ killed cells only when it was fused to RecQ^WH^ (Fig S2F).

**Figure 6.**
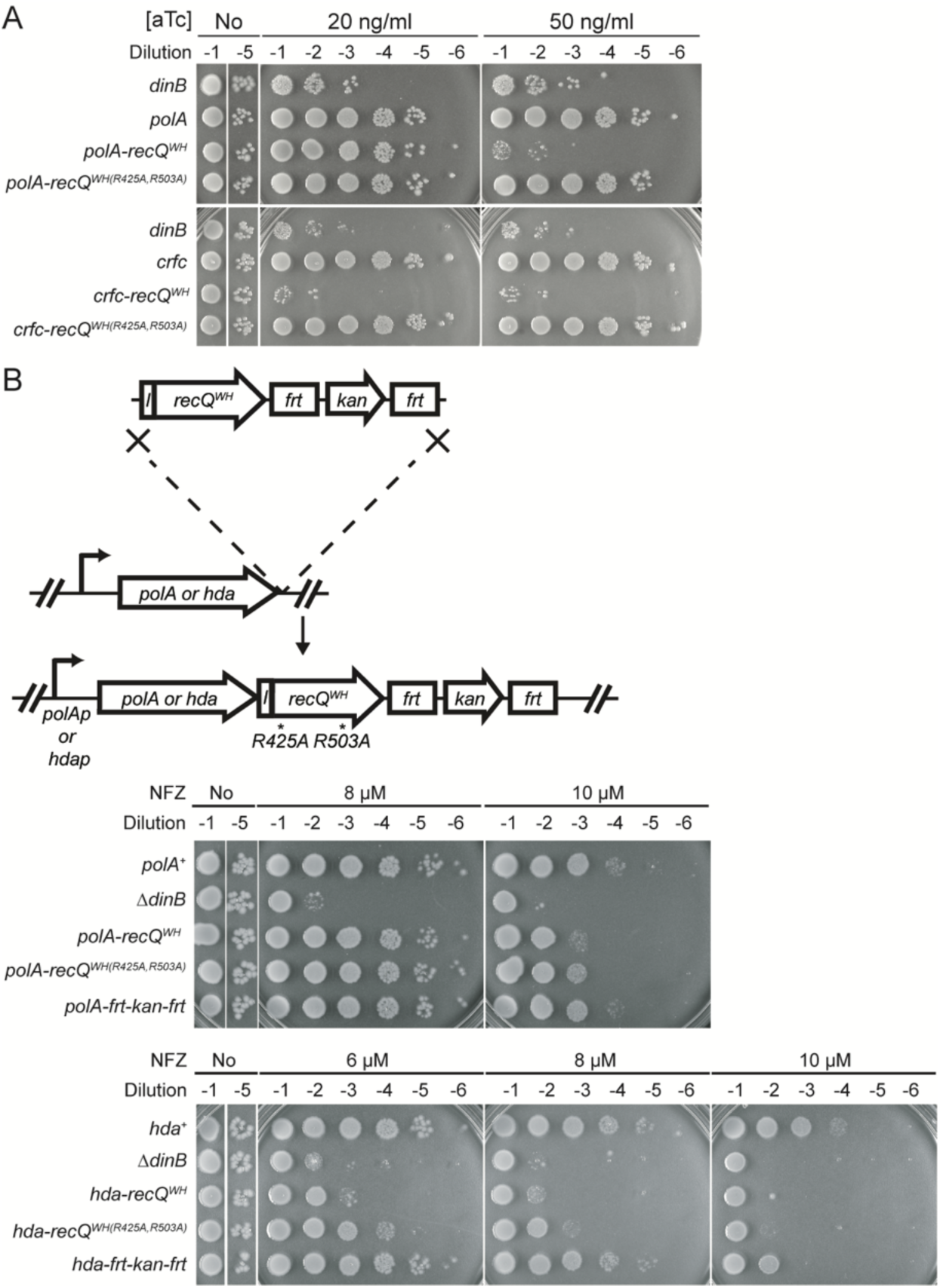
Artificial fork localization of Pol I and Hda interferes with Pol IV-mediated TLS. A. Overproduction of Pol I-RecQ^WH^ or Crfc-RecQ^WH^ kills cells. Indicated genes were expressed from a genomic tet-inducible cassette described in Fig 4G and Fig S4E. Cultures of indicated strains were serially diluted and spotted on LB-agar plates containing varying concentrations of aTc. B. Appending a RecQ^WH^ domain to the C-terminal end of Pol I or Hda sensitizes cells to NFZ. (Top) The coding sequence for *linker-recQ^WH^-frt-kan-frt* was inserted after the last coding sequence of the *polA* or the *hda* gene in the *E. coli* genome. *l*, linker; *recQ^WH^*, RecQ^WH^ domain of *E. coli* RecQ. (Bottom) Sensitivity to NFZ. Cultures of indicated strains were serially diluted and spotted on LB agar plates containing varying concentrations of NFZ.

We next asked whether artificially localizing CLIPs to the replisome through appending the RecQ^WH^ domain would interfere with Pol IV-mediated TLS. Indeed, Pol I-RecQ^WH^ cells became modestly sensitized to NFZ (Fig 6B). As the increased sensitivity to NFZ of the *polA-recQ^WH^* strain was epistatic to *ΔdinB* (Fig S6), this increased sensitivity is likely due to inhibition of Pol IV-mediated TLS by Pol I-RecQ^WH^. Moreover, mutating SSB interacting residues within the RecQ^WH^ domain (*polA-recQ^WH(R425A,R503A)^*) substantially reduced the sensitization, suggesting that inhibition of Pol IV-mediated TLS resulted from forced enrichment of Pol I near stalled replisomes. Among other CLIPs without SSB-binding activity that we examined in the same way (Crfc, Hda and LigA), we observed similar effects with Hda (Fig 6B and S6). These results suggest that CLIPs lacking SSB-binding activity are largely excluded from the replication fork.

## Discussion

As TLS polymerases are error-prone it is critical to restrict their activity to only when they are needed. How TLS polymerases are excluded from processive replisomes(Thrall et al., 2017), yet recruited to lesion-stalled replisomes has remained unclear. Previously, we showed that the ε subunit of Pol III acts as a molecular gate to limit the access of Pol IV, and presumably other CLIPs, to the β_2_ clamp(Chang et al., 2019). Here, we demonstrate that replisome-associated SSB mediates enrichment of Pol IV near replisomes upon lesion stalling, which is required for Pol IV to overcome this competitive inhibition by Pol III and mediate TLS.

### Switching between replication- and repair-competent SSB condensates at the replication fork

Our single molecule PALM imaging scheme selectively detects static Pol IV molecules around the replication fork via either direct association with DNA or interactions with replisome components. We demonstrated that Pol IV is highly enriched near lesion-stalled replisomes and nearly the entirety of static Pol IV molecules requires interaction of Pol IV with replisome-associated SSB. Intriguingly, SSB is always present on the lagging strand, yet Pol IV is not enriched at moving replisomes(Thrall et al., 2017). The local concentration of Pol IV near a replisome is determined by the net exchange of Pol IV between replisome-associated and cytosolic pools (Fig 7A). Influx into the replisome-associated pool is likely controlled by the concentration of Pol IV, which is elevated during the SOS response. However, outflux is primarily determined by the interaction of Pol IV with SSB (and possibly other replisome components). Given Pol IV is constitutively elevated in our imaging strain, the increase in static Pol IV molecules near lesion-stalled replisomes is primarily due to decreased outflux of Pol IV from the replication fork. During replication a steady state level of SSB is present on the lagging strand, forming a condensate of SSB-Ct(Harami et al., 2019), yet individual SSB molecules are rapidly turned over because the complete synthesis of an Okazaki fragment takes only ∼2 seconds, limiting the average lifetime of lagging-strand SSB molecules to around 1 second(Ogawa and Okazaki, 2002; Wu et al., 1992). This timescale may be too short to allow for association of Pol IV with SSB. Therefore, stable accumulation of Pol IV near the lesion-stalled replisome may require: 1) stabilization of SSB, 2) an increase in the amount of replisome-associated SSB and 3) potentially changes in the mode of interaction between SSB and ssDNA, all of which may be causally related.

**Figure 7.**
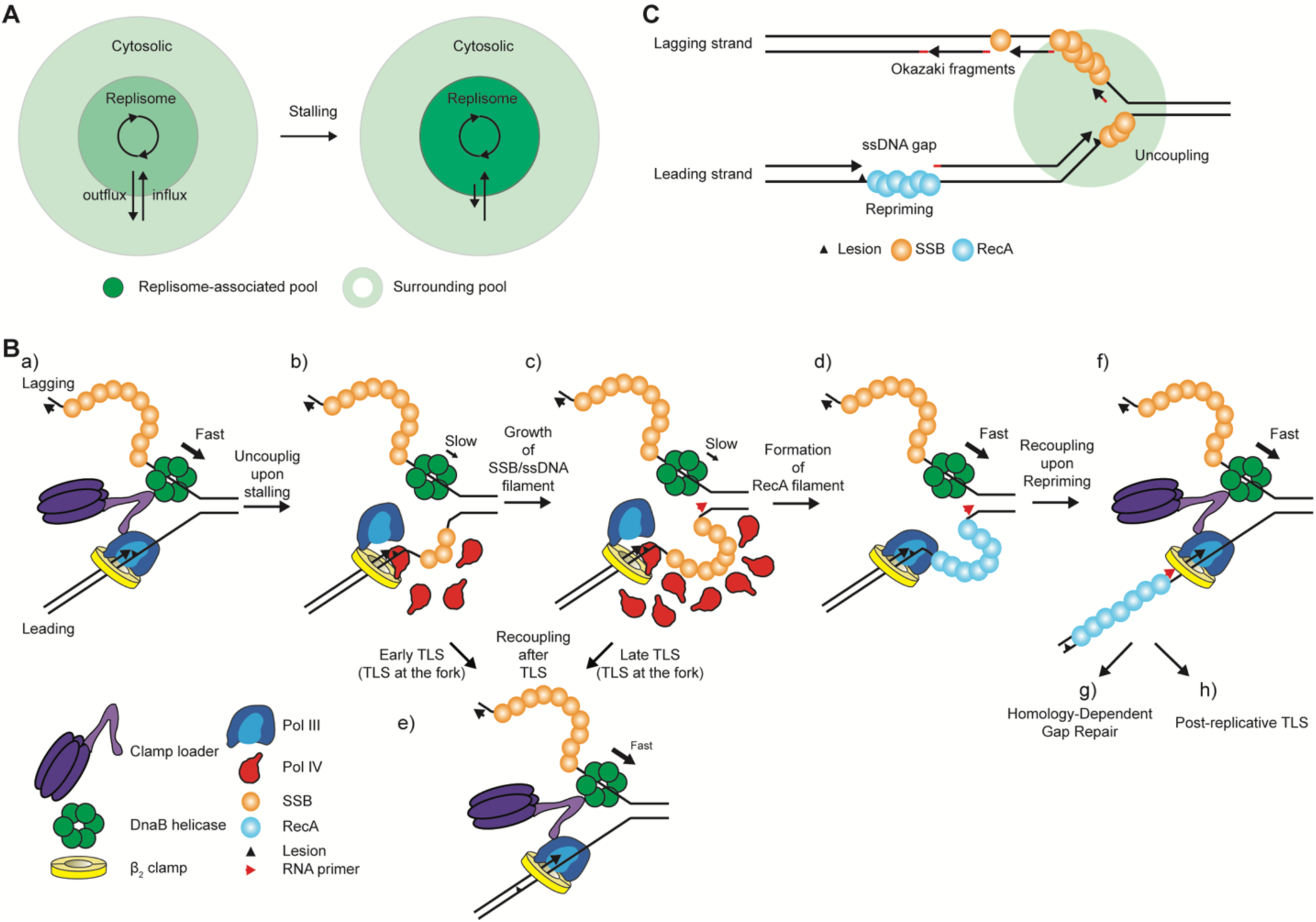
Roles of SSB in resolution pathway choice of lesion-stalled replisomes. A. Pol IV resides in either a replisome-associated or cytosolic pool. Association with SSB leads to an enrichment of Pol IV near the replisome upon fork stalling. B. Lesion stalling of the leading strand polymerase leads to uncoupling of the helicase and polymerase and the production of leading-strand ssDNA. This ssDNA gap initiates a series of SSB-dependent downstream events including TLS at the fork and RecA-mediated HDGR. C. Compartmentalization of a replication fork by SSB. A circle surrounding the replication fork represents a space where Pol IV, and possibly other SIPs, is enriched upon replication stalling at a leading strand lesion.

### Potential role of ssDNA at the replication fork in resolution pathway choice – leading vs lagging

Upon stalling of the leading strand polymerase at a lesion, the helicase becomes uncoupled from the polymerase and slows from an unwinding rate of ∼1000 bp/sec to a rate of ∼30 bp/sec, the intrinsic unwinding rate of DnaB(Kim et al., 1996) (a to b Fig 7B). This slow unwinding generates a growing stretch of ssDNA on the leading strand template that continues until leading strand synthesis is resumed by either TLS at the fork or DNA synthesis is reprimed downstream of the blocking lesion(Yeeles and Marians, 2013). SSB molecules likely associate with this leading-strand ssDNA (gap) and interact with Pol IV, enriching Pol IV near stalled replisomes (b and c in Fig 7B).

Growth rates and length distributions of leading-strand ssDNA gaps in cells have not been directly measured but are determined by both the unwinding rate of the DnaB helicase and the rate of repriming leading strand synthesis. Both *in vivo* and *in vitro* observations estimate that the leading-strand ssDNA gap is roughly a few hundred nucleotides(Rupp and Howard-Flanders, 1968; Yeeles and Marians, 2013), which can bind several SSB molecules(Lohman and Overman, 1985). Unlike SSB on the lesion-free lagging strand template, which is rapidly displaced by the lagging strand polymerase, SSB molecules on the leading-strand ssDNA are likely much more stable as DNA synthesis is temporarily blocked by the lesion. The lifetime of leading-strand SSB is limited by how quickly stalling is resolved by TLS because upon TLS Pol III resumes rapid synthesis and displaces SSB (b or c to e in Fig 7B). The rate of TLS over a lesion is determined by two factors; 1) cellular levels of cognate TLS polymerases and 2) inherent efficiency of TLS over a specific lesion by the cognate TLS polymerase.

Shortly after stalling of the leading strand polymerase at a lesion, a relatively short stretch of ssDNA (35 or 65 nts) is generated which can bind one SSB_4_ molecule. SSB_4_ in this and similarly short SSB_4_/ssDNA complexes interacts with Pol IV, elevating the local Pol IV concentration. This relatively low enrichment of Pol IV may be sufficient for efficient TLS over blocking lesions that can be easily replicated past, such as N^2^-FFdG. Rapid TLS (TLS at the fork) leads to recoupling with the helicase (b to e in Fig 7B) and prevents the leading-strand ssDNA from growing long enough to allow for formation of a RecA/ssDNA nucleofilament and/or repriming.

Alternatively, when stalling persists at strongly blocking lesions on the leading strand, such as 3meA and benzopyrene (BaP), a growing tract of SSB/ssDNA recruits more Pol IV, further elevating its local concentration (c to e in Fig 7B). This relatively high enrichment of Pol IV may facilitate TLS over strong blocks (late TLS). Failure to carry out TLS leads to displacement of SSB by RecA(Bell et al., 2012; Morimatsu and Kowalczykowski, 2003) and a concomitant drop in Pol IV local concentration (c and d in Fig 7B). Persistent stalling also triggers repriming (d to f in Fig 7B), which leads to a ssDNA gap. These gaps are likely filled in by Homology-Dependent Gap Repair (HDGR) due to the strong recombinogenic activity of the RecA/ssDNA filament (f to g in Fig 7B) while only a small fraction is filled in by TLS in a post-replicative manner(Fuchs, 2016) (f and h in Fig 7B). This minor utilization of TLS at the gap is also partly due to weak enrichment of Pol IV near the gap, which is insufficient to overcome the ε kinetic barrier(Chang et al., 2019). Consistent with this model, the majority of BaP-stalled replisomes are resolved by HDGR(Chang et al., 2019) whereas N^2^-FFdG-stalled replisomes are predominantly resolved by TLS (Fig 5G), Similar to our observations on the leading strand, weakening the Pol IV-SSB interaction also reduced the utilization of TLS at a lagging strand lesion (Fig 5D), suggestive of a common regulatory mechanism on both strands.

### A possible mechanism underlying the stalling-dependent interaction between SSB and Pol IV

The salient feature of the Pol IV-SSB interaction is that it requires association of SSB with ssDNA. Given that the interaction between Pol IV and SSB is largely mediated through SSB-Ct, it is conceivable that SSB-Ct is only accessible to Pol IV when SSB is bound to ssDNA. Indeed, SSB-Ct in bare SSB is in equilibrium between a “bound” state in which SSB-Ct is intramolecularly associated with the OB domain of SSB and an “unbound” state free of intramolecular interaction. As the SSB-Ct binding surface and the ssDNA binding surface within the OB domain of SSB are overlapping (Raghunathan et al., 2000; Shishmarev et al., 2014), it is possible that association of ssDNA with SSB competitively displaces bound SSB-Ct, making it competent to interact with SIPs.

Unlike Pol IV, we found that PriA and Exonuclease I could interact with bare SSB and association of SSB with ssDNA only increased fractional binding without significantly changing the binding affinity (Fig S1D). These observations imply that a measurable fraction of SSB-Ct in bare SSB is in the unbound state and association of ssDNA with SSB merely increases the fraction. Therefore, it is unclear how ssDNA increases both fractional binding and the binding affinity for Pol IV. Given that Pol IV has a weak ssDNA binding activity, it is possible that ssDNA cooperatively promotes the interaction between Pol IV and SSB along with increasing the fraction of unbound SSB-Ct.

### Resolution pathway choice and mutagenesis

Resolution pathway choice of lesion-stalled replisomes determines not only tolerance of bacterial cells to damaging agents but also the extent of damage-induced mutagenesis. TLS at the fork enables a replisome to directly replicate past a blocking lesion without creating a ssDNA gap and is likely limited to the vicinity around the lesion(Chang et al., 2019). Moreover, errors made by TLS polymerases may be corrected by the proofreading activity of the replicative polymerase, which switches back to resume processive replication(Fujii and Fuchs, 2004; Jarosz et al., 2009). In contrast, repriming of DNA replication downstream of the lesion, leaves a long ssDNA gap (>200 nucleotides) that is repaired primarily by a high-fidelity recombination-dependent gap filling mechanism. However, a small fraction of these gaps are filled in by the combined actions of replicative and TLS polymerases in a highly mutagenic manner (TLS at the gap) (Isogawa et al., 2018). This widespread mutagenesis that results during TLS at the gap is more likely to lead to functional genetic alterations compared with the localized mutagenesis that occurs adjacent to the lesion during TLS at the fork. In this study, we demonstrate that SSB facilitates TLS at the fork, resulting in the suppression of repriming. Moreover, consistent with TLS at the fork being less mutagenic than TLS at the gap, MMS-induced mutagenesis is highly elevated in a *dinB^T120P^* strain compared with the wild-type *dinB*(Scotland et al., 2015).

### Spatial segregation of CLIPs

Given that all CLIPs interact with the β_2_ clamp via a common binding site, it is conceivable that competitive clamp binding among CLIPs would interfere with the action of each other. Notably, TLS polymerases also interact with SSB whereas other CLIPs do not, and we demonstrate that Pol IV is highly enriched near lesion-stalled replisomes through the interaction with replisome-associated SSB. Therefore, SSB binding activity serves as a fork-localization signal and thus CLIPs lacking SSB-binding activity are likely excluded from the replication fork even upon replisome stalling. Interference among CLIPs at the fork may be minimized by this spatial segregation. Consistent with this model, Pol I, despite its high cellular abundance, only interferes with Pol IV mediating TLS at the fork when artificially localized to replication forks by the SSB-interacting RecQ^WH^ domain.

Conversely, actions of Pol I and LigA at the junction between a completed Okazaki fragment and a downstream RNA primer on the lagging strand template are likely not inhibited by CLIPs with SSB binding activity. As the ssDNA gaps between immediate RNA primers on the lagging strand template are filled in by Pol III during Okazaki fragment synthesis, lagging-strand SSB molecules are rapidly displaced and the DNA/RNA junction, which is ligated by sequential actions of Pol I and LigA, becomes spatially separated from the fork by a long rigid stretch of dsDNA (Fig 7C). This loss of SSB-coated DNA ensures that CLIPs with SSB binding activity, such as Pol IV, are barely enriched near the DNA/RNA junction. We propose, that under cellular concentrations, CLIPs require local enrichment to gain access to the β_2_ clamp. This hierarchical recruitment is mediated by interactions with other factors, such as SSB, or DNA structures that act to locally segregate CLIPs to their site of action.

## Acknowledgements

We thank Kelly Arnett (Center for Macromolecular Interactions, Harvard Medical School) for training and assistance in using a CD spectropolarimeter, James Keck (University of Wisconsin, Madison) for providing the fluorescein-conjugated SSB-Ct peptides and Deyu Li (University of Rhode Island) for providing the N^2^-FFdG-containing oligomer. This work was supported by National Institutes of Health grants R01 GM114065 (to J.J.L), F32 GM113516 (to E.S.T),

Authors declare no conflicts of interest

## Author contributions

S.C., E.S.T., L.L., V.P. and J.J.L. designed and performed research. S.C., E.S.T., L.L., V.P. and J.J.L. analyzed data. S.C., E.S.T., L.L., V.P. and J.J.L. wrote the paper.

---

**Supplementary Figure 1.**
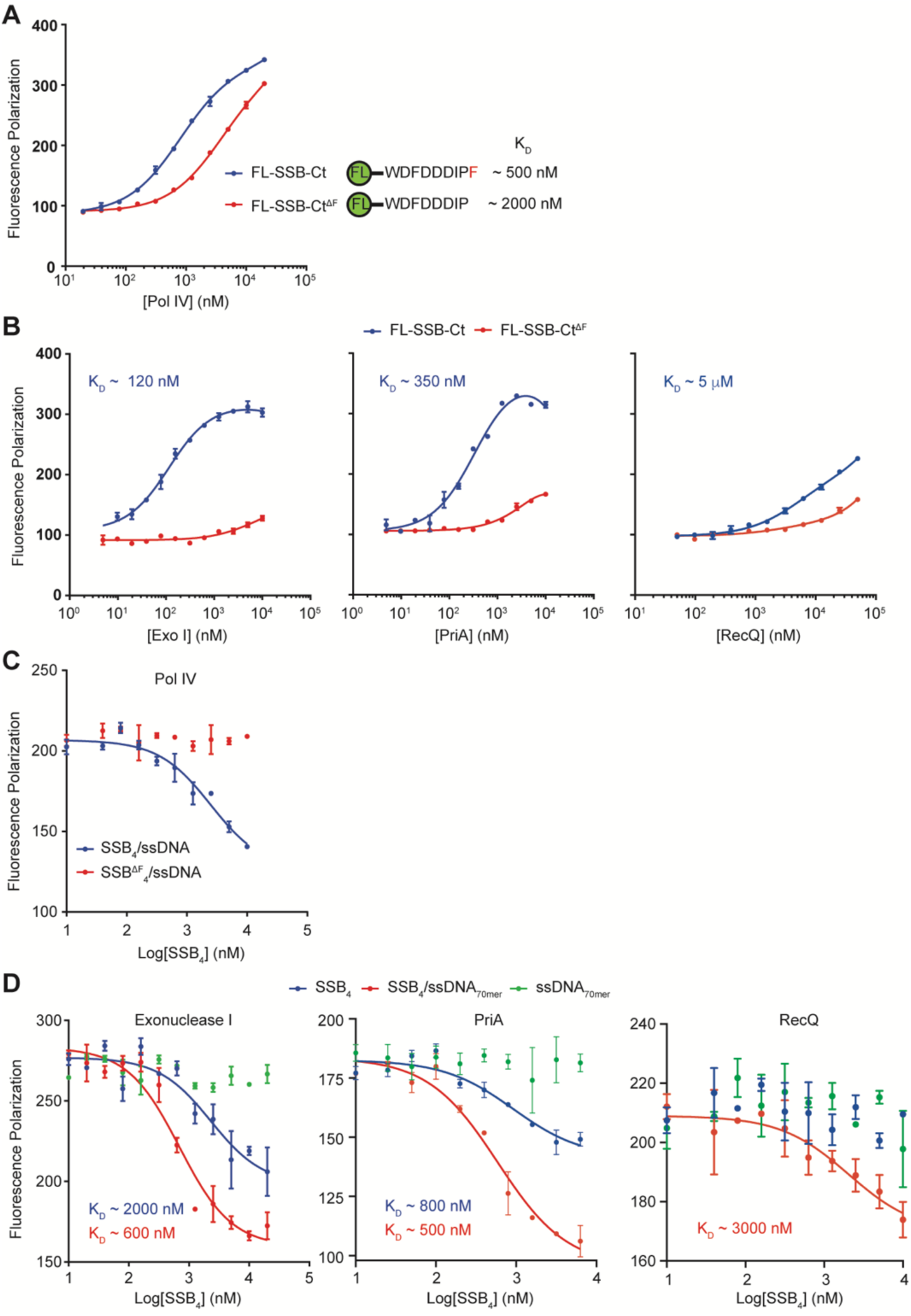
A. Interaction between purified Pol IV and various FL-SSB-Ct peptides as measured by FP. B. Interaction of 1) Exo I, 2) PriA and 3) RecQ with either FL-SSB-Ct or FL-SSB-Ct^ΔF^. C. Ultimate phenylalanine of SSB is required for interaction with Pol IV. Competitive displacement of bound FL-SSB-Ct from Pol IV by indicated SSB. D. Competitive displacement of bound FL-SSB-Ct by full-length SSB from 1) Exonuclease I, 2) PriA and 3) RecQ.

**Supplementary Figure 2.**
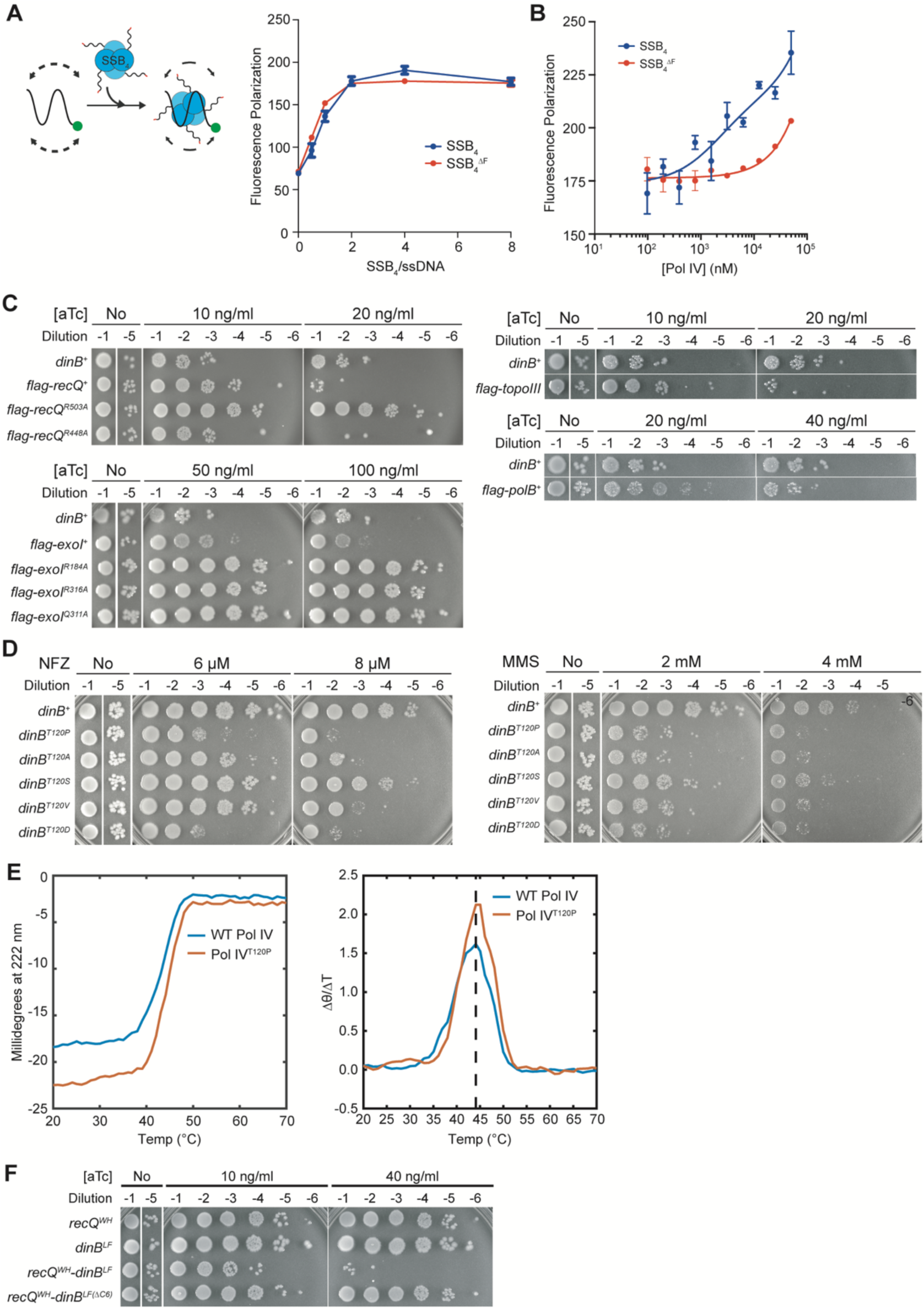
A. Full-length SSB and SSB^ΔF^ form nucleoprotein complexes with T_71_-FAM. (Left) Binding scheme. (Right) Interaction T71-FAM with either 1) SSB or 2) SSB^ΔF^ measured by FP. B. Deleting the ultimate phenylalanine of full-length SSB abolishes interaction with Pol IV. C. Intracellular overproduction of 1) RecQ, 2) Exonuclease I, 3) Topoisomerase III or 4) Pol II leads to massive cell death. Cell death by RecQ or Exonuclease I depends on interaction with SSB; indicated *recQ* and *exoI* mutations diminish their interaction with SSB. D. Overproduction of RecQ^WH^-Pol IV^LF^ kills cells while overproduction of neither RecQ^WH^ nor Pol IV^LF^ alone is sufficient for cell killing. E. Pol IV^T120P^ retains wild-type thermal stability. (Left) Changes in ellipticity of Pol IV and Pol IV^T120P^ at 222 nm upon increasing temperature. (Right) 1^st^ derivative of the unfolding curves shown in the left panel. The maxima correspond to the midpoints (T_MS_) of the unfolding curves. F. Mutating Thr^120^ of *dinB* to various amino acids sensitizes cells to NFZ and MMS.

**Supplementary Figure 3.**
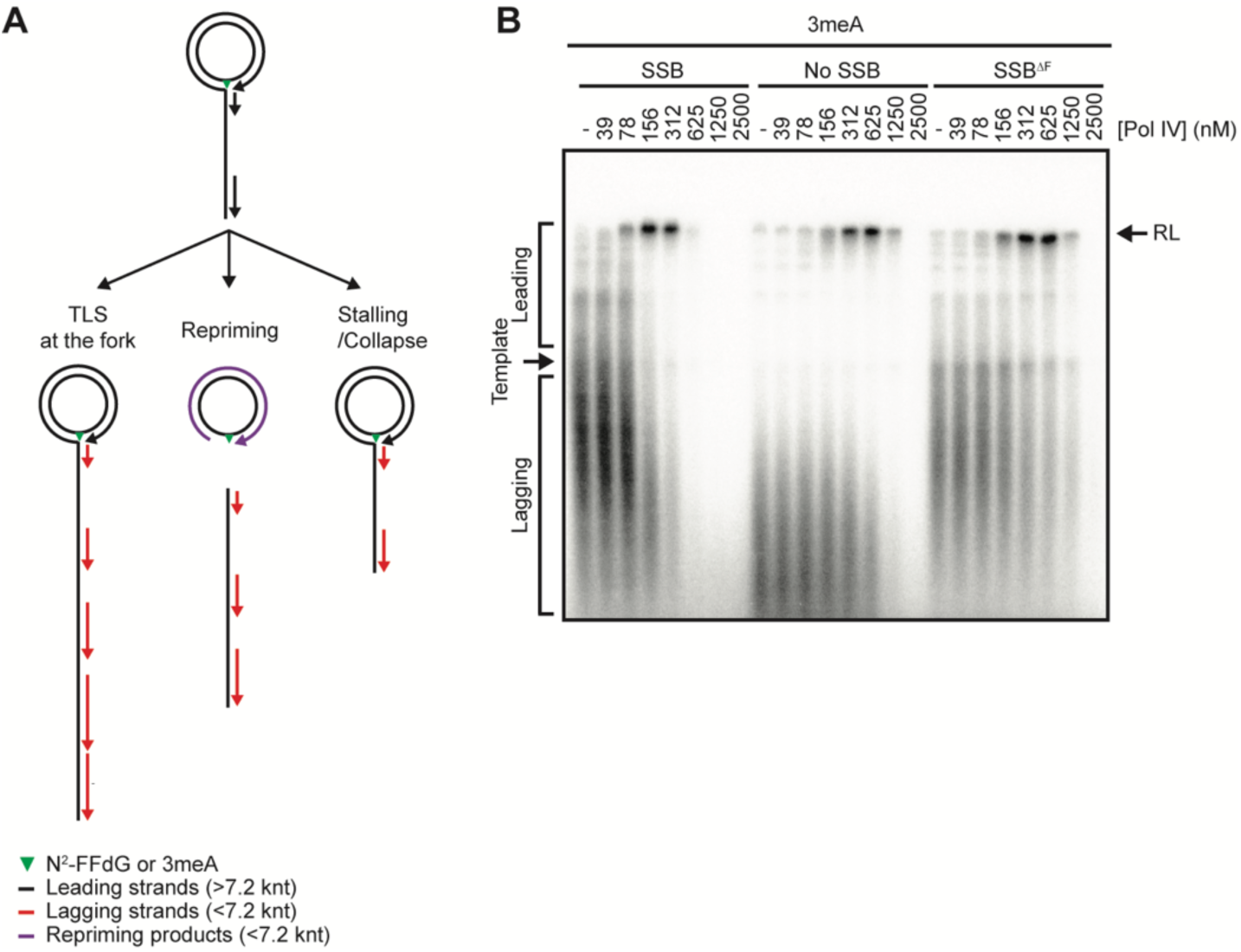
A. Fates of stalled replication and associated replication products in our rolling circle replication-based assay. B. Interaction of SSB with Pol IV potentiates Pol IV-mediated TLS over 3meA. Only the resolution-limited leading strand bands are presented in Fig 3D.

**Supplementary Figure 4.**
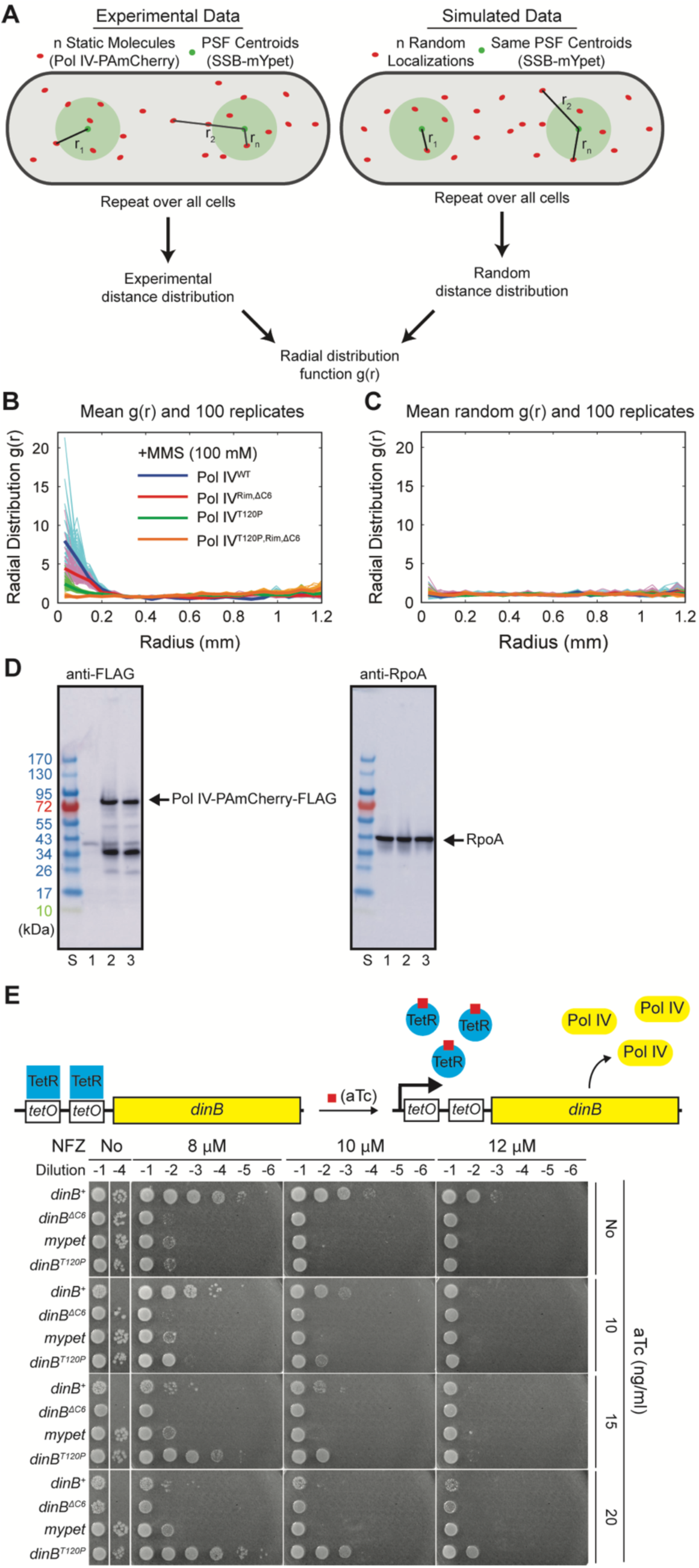
A. Cartoon of radial distribution function analysis. (Left) Experimental distances between static Pol IV-PAmCherry molecules and the nearest SSB-mYPet foci are calculated. (Right) For each cell, the same number of random localizations are generated and the distances to the same SSB-mYPet foci are determined. This calculation is performed over all cells and the resulting experimental distribution is normalized by the random distribution to give the radial distribution function *g(r)*. This procedure is repeated 100 times to generate 100 *g(r)* curves, which are averaged to give the final mean *g(r)* curve. B. Variability of calculated Pol IV-SSB radial distribution function curves for Pol IV^WT^ and mutants in cells treated with 100 mM MMS. The 100 *g(r)* curves (thin lines) and mean *g(r)* curves (thick lines) are plotted for each data set. C. As in B, but for the random *g(r)* curves. D. Comparable expression of Pol IV-PAmCherry-FLAG and Pol IV^T120P^-PAmCherry-FLAG in the imaging strains. FLAG-tagged proteins were detected by Western blotting with an anti-FLAG antibody. S. protein size markers; 1, the parent imaging strain; 2, an imaging strain expressing Pol IV-PAmCherry-FLAG; 3, an imaging strain expressing Pol IVT^120P^-PAmCherry-FLAG. RpoA, a loading control, was probed by Western blotting with an anti-RpoA antibody. E. (Top) A tet-inducible ectopic expression cassette engineered within the *lamB* gene in a *ΔdinB* strain. (Bottom) Normalization of tolerance to NFZ by induced expression of indicated genes.

**Supplementary Figure 5.**
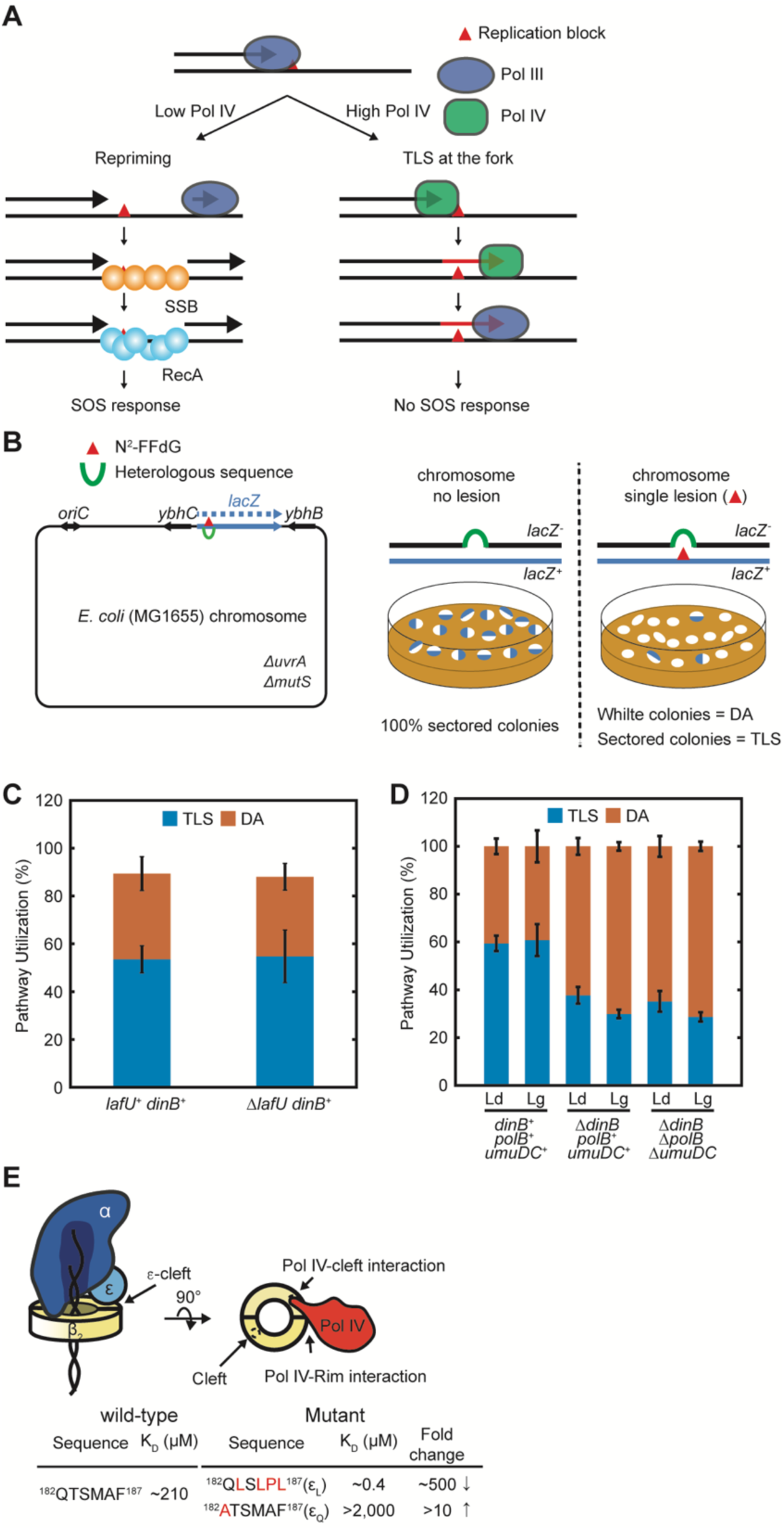
A. Repriming by lesion-stalled replisomes creates ssDNA gaps, which eventually leads to induction of the SOS response. B. (Left) Site-specific insertion of a single N^2^-FFdG adduct into the *E. coli* genome in either the leading or lagging strand templates. (Right) Resolution of lesion-stalled replication through TLS leads to formation of blue-sectored colonies. White colonies represent resolution through DA. C. Deletion of *lafU*, which is replaced with a *frt-kan-frt* cassettes as a *dinB*-linked marker for P1 transduction, does not influence Pol IV-mediated TLS over N^2^-FFdG in cells. D. TLS over N^2^-FFdG in cells is primarily mediated by Pol IV. Deleting *dinB* reduces resolution of N^2^-FFdG-stalled replisomes through TLS from ∼60 to ∼30%. Additional deletion of both *polB* and *umuDC* does not further reduce resolution through TLS. The residual TLS (∼30%) in the *ΔdinB ΔpolB ΔumuDC* background is presumably mediated by Pol III. However, as Pol IV outcompetes Pol III in mediating TLS over N^2^-FFdG, the actual contribution of Pol III to TLS in the *dinB^+^* background is likely less than estimated in the *ΔdinB* background. Ld, leading strand lesion; Lg, lagging strand lesion. E. Clamp binding mutations of *dnaQ*. *dnaQ(ε_Q_)*, weakening mutation; *danQ(ε_L_)*, strengthening mutation.

**Supplementary Figure 6.**
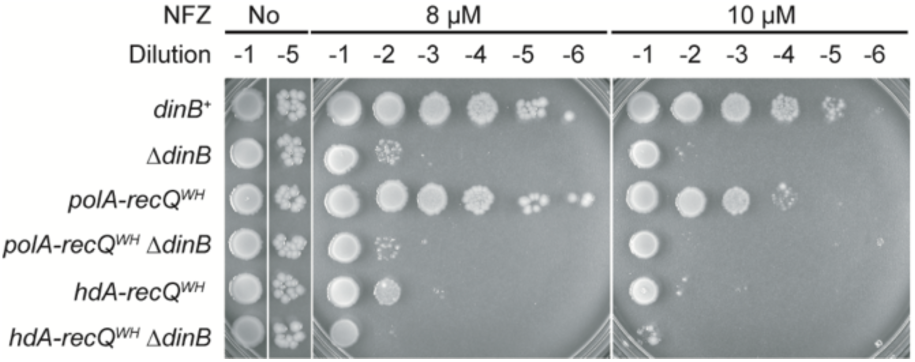
Sensitivity of the *polA-recQ^WH^* and the *hda-recQ^WH^* strains to NFZ is epistatic to *ΔdinB*. Cultures of indicated strains were serially diluted and spotted on LB agar plates containing varying concentrations of NFZ.

## Materials and Methods

### Materials

#### Purified proteins

Purified αεθ, αε_Q_θ, αε_L_θ, α, DnaB, clamp loader complex (τ_3_δδ’χψ), β_2_ clamp, DnaG and SSB were purified as previously described(Chang et al., 2019). Wild-type Pol IV and its variants were expressed in BL21(DE3) pLysS lacking the native *dinB* and purified as previously described(Jarosz et al., 2006). PK-His6-Pol IV^LF^ was purified using Ni-NTA resin followed by size exclusion chromatography with a Superdex100 10/00 column.

#### Reagents

FL-SSB-Ct (FAM-WMDFDDDIPF), FL-SSB-Ct^DF^ (FAM-WMDFDDDIP) and FL-SSB-Ct^113^ (FAM-WMDFDDDISF) were provided by James Keck (Univ. of Wisconsin, Madison) (Lu et al., 2009). N^2^-furfuryl dG-containing oligomer (5’-CTACCT/N^2^-furfuryl-dG/TGGACGGCTGCGA-3’) were provided by Deyu Li (Univ. of Rhode Island). 3-deaza-methyl dA-containing (GCTCGTCAGACG/3-deaza-3-methylA/TTTAGAGTCTGCAGTG) was synthesized by IDT (IA, USA). 3′ FAM-conjugated ssDNA of 71 thymidine (T_71_-FAM) was synthesized by IDT (IA, USA).

#### Chemicals

Methyl methanesulfonate (MMS, Sigma, 129925), Nitrofurazone (NFZ, Fluka, PHR1196), Anhydrotetracycline (aTc, TaKaRa)

#### Fluorescence polarization (FP)-based binding assays

Binding assays were performed in binding buffer (50 mM Hepes-HCl pH7.5, 100 or 200 mM NaCl, 0.05% Tween20, 10 mM 2-mercaptoethanol; BB100 or 200, binding buffer containing 100 or 200 mM NaCl). FL-SSB-Ct or its variants, or T_71_-FAM was incubated with various SSB interacting proteins and SSB at room temperature (RT), and fluorescence polarization of fluorescein was measured at 25 °C using a SpectraMax M5 (Molecular Devices).

For direct binding assays with fluorescein-conjugated SSB-Ct peptides, a labeled peptide (20 nM) was incubated with a binding partner at varying concentrations in BB100 at RT. For direct binding assays with fluorescein-conjugated ssDNA (T_71_-FAM), preformed SSB/T_71_-FAM (50 nM) was incubated with a binding partner at varying concentrations in BB200 at RT. To preform SSB/T_71_-FAM, 1.2:1 ratio of SSB (or SSB^ΔF^) and T_71_-FAM was incubated in BB200 at RT before incubated with a binding partner (Fig S2A). The equilibrium dissociation constant of a labeled peptide was determined by fitting the following model to measured FP values, F = Fmax X C/(K_D_ + C) + NS X C + Background, wherein F is FP, Fmax is the maximum FP, C is the concentration of a SIP, K_D_ is the equilibrium dissociation constant, NS is the slope of nonspecific binding and Background is the amount of nonspecific binding.

For competition binding assays with unlabeled peptides, purified Pol IV was incubated with FL-SSB-Ct peptide (20 nM) and unlabeled peptides at varying concentrations in BB100. For competition binding assays with either bare or ssDNA-wrapped full-length SSB (or SSB*^Δ^*^F^), a fixed amount of a binding partner protein was incubated with FL-SSB-Ct peptide (20 nM) and either bare SSB or preformed SSB/ssDNA at varying concentrations in BB200 at RT. To preform ssDNA-wrapped full-length SSB, ssDNA and SSB (or SSB*^Δ^*^F^) were mixed and incubated in BB200 at RT before titrated into binding reactions. To obtain IC50, the following model was fitted to measured FP values, F = Fbottom + (Ftop – Fbottom)/(1 + 10^(C-logIC50)^), wherein Ftop and Fbottom are plateaus in FP, C is the concentration of an unlabeled competitor, and IC50 is the concentration of a competitor that displaces 50% of bound FL-SSB-Ct. Then the IC50 was used to calculate the equilibrium dissociation constant of an unlabeled competitor by logIC50 = log(10^logKi^ X (1 + [FL-SSB-Ct]/K_D_^FL-SSB-Ct^)), wherein Ki is the equilibrium dissociation constant of an unlabeled competitor in M, [FL-SSB-Ct] is concentration of FL-SSB-Ct in nM and K_D_^FL-SSB-Ct^ is the equilibrium dissociation constant of FL-SSB-Ct in nM for the binding partner.

#### Measuring the damage-induced SOS-response

The DNA damage-induced SOS response was measured as previously described(Chang et al., 2019). Briefly, SOS reporter strains were cultured in Luria Broth (LB) at 37 °C with aeration until OD_600_ reached ∼0.3. Cultures were treated with MMS (7 mM final) and further incubated at 37 °C with aeration for an additional 1 hour. Then cells were fixed with formaldehyde (4 %), thoroughly washed with phosphate-buffered saline (PBS, pH 7.4) and finally resuspended in PBS. GFP fluorescence from individual cells was measured by flow cytometry using an Accuri C6 (BD Biosciences).

#### Measuring sensitivity to NFZ, MMS and overproduced proteins

Overnight cultures were diluted in 0.9 % NaCl to OD_600_ = 1.0 and serially diluted by 10-fold to 10^6^ -fold. Serially diluted cultures were spotted on LB-agar containing NFZ, MMS or anhydrotetracycline. Plates were photographed after a 15 hour incubation at 37 °C.

#### E. coli strains

Refer to the supplementary table 1 for *E. coli* strains used in this study.

### Construction of *E. coli* strains

#### General approach

Strains were constructed as previously described(Chang et al., 2019). Briefly, mutations of interest including insertions were introduced into the *E. coli* genome by lambda Red recombinase-mediated allelic exchange(Sharan et al., 2009). frt-flanked antibiotic selection markers were used to either disrupt a gene or to create a linkage to an allele of interest for P1 transduction.

#### Strains bearing a tetracycline-inducible (ptet) expression cassette

The Z2 locus bearing expression cassettes for *tetR* and *lacIq*(Lutz and Bujard, 1997) was transferred to MG1655 by P1 transduction. In the resulting strain (MG1655 Z2), a tetracycline inducible expression cassette (tccctatcagtgatagagattgacatccctatcagtgatagagatactgagcactactagagaaagaggagaaatactag **ATGGACTACAAAGACGATGACGACAAG**gaattctagtgctagtgtagatcgctactagagccaggcatcaaataaaacgaaag gctcagtcgaaagactgggcctttcgttttatctgttgtttgtcggtgaacgctctctactagag; two tetO sites (red), a RBS (green), a FLAG epitope coding sequence (bold black), EcoR I site (orange), T1 terminator of the *E. coli rrnB* (blue)) followed by *frt-kan-frt* was inserted between 621 and 658 nucleotides of *lamB* by lambda Red recombinase-mediated allelic exchange, generating MG1655 *ΔlamB::ptet-frt-kan-frt*. To insert a coding sequence to be induced into the expression cassette, the kan cassette was first flipped out by transiently expressing flippase in MG1655 *ΔlamB::ptet-frt-kan-frt*, resulting in MG1655 *ΔlamB::ptet-frt*. Then, the remaining *frt* and flanking elements were replaced with the coding sequence of interest either with N-terminal flag or without flag followed by *frt-kan-frt* through lambda Red recombinase-mediated allelic exchange. Tet-inducible expression strains created in this way lack T1 terminator. Various alleles of interest were also introduced into the inducible strains by P1 transduction.

#### Making endogenous fusions to RecQ^WH^ domain of the *E. coli recQ*

The sequence for *linker-recQ^WH^-frt-kan-frt* was inserted in frame to the 3’ end of the last coding sequence of a host gene in the *E. coli* genome by lambda Red recombinase-mediated allelic exchange (Fig 6B). Linker, SAGSAAGSGEF(Uphoff et al., 2013); RecQ^WH^, amino acid number 408 to 523 of RecQ.

### Rolling circle replication

#### Construction of control and lesion-containing rolling circle DNA templates

Rolling circle templates were constructed as previously described(Chang et al., 2019). Briefly, a lesion-containing oligomer (N^2^-furfuryl dG, 5’-CTACCT/N^2^-furfuryl dG/TGGACGGCTGCGA-3’; 3-deaza-methyl dA, GCTCGTCAGACG/3-deaza-3-methylA/TTTAGAGTCTGCAGTG) (Benson et al., 2014; Jarosz et al., 2009) was ligated into EcoRI-linearized M13mp7L2, generating a lesion-containing closed circular ssDNA. This circular ssDNA was converted into a 5’-tailed dsDNA by T7 DNA polymerase-catalyzed extension of a primer (N^2^-furfuryl dG, T_36_GAATTTCGCAGCCGTCCACAGGTAGCACTGAATCATG; 3-deaza-methyl dA, T_36_-TTCACTGCAGACTCTAAATCGTCTGACGAGCCACTGA) that was annealed over the lesion-containing region of the circular ssDNA. Control templates were constructed in the same way but with lesion-free oligomers of the same sequences.

#### Rolling circle replication

Rolling circle replication was performed as previously described(Chang et al., 2019) (Fig 3A). Briefly, the *E. coli* replisome was reconstituted on either a control or a lesion-containing rolling circle template with purified replisome components. To assemble replisomes, a mixture of a rolling circle template (1 nM), τ_3_δδ’χψ (20 nM), Pol III core (20 nM), β_2_ clamp (20 nM), DnaB_6_ (50 nM), ATPγS (50 μM) and dCTP/dGTP (60 μM each) was prepared in HM buffer (50 mM Hepes (pH 7.9), 12 mM Mg(OAc)2, 0.1 mg/mL BSA and 10 mM DTT) on ice, then incubated at 37 °C for 6 min. After the assembly, replication reactions were initiated by adding a 10X initiation mixture (1 mM ATP, 250 μM each CTP/GTP/UTP, 60 μM each dATP/dTTP, [α-^32^P]-dATP, 200 μM SSB_4_, 100 nM DnaG). After incubation at 37 °C for 12 min, replication reactions were quenched by adding EDTA (25 mM final). All the concentrations are final concentrations in the replication reactions.

Replication products were separated in a 0.6 % alkaline denaturing alkaline agarose gel and visualized by autoradiography. Radioactive signals were quantitated using ImageJ. Leading strand synthesis was quantified by integrating signals above the template. These values for the lesion-free control template and the lesion-containing template were defined “processive replication” and “TLS” respectively. Relative band intensities were calculated with respect to the band intensity in the absence of Pol IV for processive replication and the maximal band intensity in the presence of Pol IV for TLS, respectively. In some figure panels, only resolution-limited bands were presented for easy comparison among various conditions.

#### Detecting repriming products by Southern blot

Repriming products were detected by Southern blotting as previously described(Chang et al., 2019). Briefly, rolling circle replication products created in the absence of [α-^32^P]-dATP were separated in a 0.6% alkaline denaturing agarose gel and transferred to a Nylon membrane (Hybond – XL, GE Healthcare) by a downward method after mildly cleaved in 0.25 M HCl. Transferred DNAs were UV cross-linked to the membrane with a UV cross-linker (UV Stratalinker 2400, Stratagene). After incubating the cross-linked membranes in a blocking buffer (ULTRAhyb, Invitrogen, AM8670) for 5 hours at 42 °C, a mixture of three 5’ [^32^P]-labeled oligomer probes shown in Fig 5A (oligomer 1, 5′-ACCGATTTAGCTTTATGCTCTGAGGCTTTATTGCTTAATT-3′; oligomer 2, 5′-AATGCTACTACTATTAGTAGAATTGATGCCACCTTTTCAG-3′; oligomer 3, 5′-TAAAGGCTTCTCCCGCAAAAGTATTACAGGGTCATAATGT-3′) were added and hybridized with the membrane for 20 hours at 42 °C with continuous agitation. Radioactive signals were detected by autoradiography and repriming products were quantitated using ImageJ.

#### Western blot

The expression levels of different Pol IV-PAmCherry fusion proteins were measured by Western blot as previously described(Thrall et al., 2017). Cell lysates of imaging strains expressing different C-terminally FLAG-tagged Pol IV-PAmCherry variants were probed with an anti-FLAG antibody or an anti-RpoA antibody for Pol IV-PAmCherry-FLAG and RpoA, a loading control, respectively.

Cultures were grown in 50 mL volumes following the same growth procedure as in imaging experiments. Cells were harvested at OD_600nm_ ≈ 0.15, resuspended in 1 mL chilled DI H_2_O, and pelleted again. The cell pellet was resuspended and lysed in B-PER Bacterial Protein Extraction Reagent (Thermo Scientific #78248) supplemented with lysozyme (EMD #5950), Benzonase nuclease (Novagen #70746) and EDTA-free protease inhibitor cocktail (Roche #04693159001).

Samples were run on a SDS-PAGE gel (Bio-Rad #4561086: 4-15 Mini-PROTEAN TGX) along with BioReagents EZ-Run Prestained Rec Protein Ladder (Fisher #BP3603), then transferred to a polyvinylidene difluoride membrane (PerkinElmer #NEF1002001PK: PolyScreen PVDF Hybridization Transfer Membrane). After blocked in TBST containing 5% skim milk, one membrane was probed with a rabbit anti-FLAG antibody raised against the antigen Ac-C(dPEG4)DYKDDDDK-OH (a gift of Johannes Walter, Harvard Medical School) and the other was probed with a mouse anti-RpoA monoclonal antibody (BioLegend #663104; Clone 4RA2, Isotype Mouse IgG1). A goat anti-rabbit IgG-HRP antibody (Jackson ImmunoResearch #111-035-003) and a rabbit anti-mouse IgG-HRP antibody (Jackson ImmunoResearch #315-035-003) were used as secondary antibodies for the anti-FLAG and the anti-RpoA blots respectively. The membranes were visualized using an Amersham Imager 600 with HyGLO chemiluminescent HRP antibody detection reagent (Denville Scientific #E2400).

#### Circular Dichroism

The protein storage buffer for purified Pol IV and Pol IV^T120P^ was exchanged to 10 mM potassium phosphate (pH 7.5) with Micro Bio-Spin P-30 Chromatography Columns (Bio-RAD) right before the measurements. Circular dichroism of the buffer-exchanged Pol IV and Pol IV^T120P^ at 1 μM was measured using a CD spectropolarimeter (Jasco J-815) with a Peltier temperature controller. To monitor thermal denaturation, dichroic activities of Pol IV and Pol IV^T120P^ at 222 nm were recorded at varying temperatures.

### In vivo TLS assays

#### In vivo TLS assay strains

All *in vivo* TLS assay strains are derivatives of strains FBG151 and FBG152(Esnault et al., 2007; Pagès et al., 2012). Various *dinB*, *polB* and *umuDC* alleles were introduced to the *in vivo* TLS assay strains by P1 transduction and selected for linked-selection markers, and then selection markers were flipped out by transient expression of flippase. The *in vivo* TLS assay strains used in this study also carry the plasmid pVP135, which allows IPTG-inducible expression of the int–xis genes for site-specific genomic integration of lesion-containing or control plasmids.

#### In vivo assay to measure TLS

In vivo assays were performed as previously described(Pagès et al., 2012). 40 μL of electrocompetent cells were transformed with 1 ng of the lesion-containing plasmid mixed with 1 ng of pVP146 (internal control for transformation efficiency) by electroporation using a GenePulser Xcell (BioRad) at 2.5 kV, 25 μF and 200 Ω. Transformed cells were first resuspended in 1 mL of super optimal broth with catabolic repressor (SOC), and then 500 μL of the cell resuspension was transferred into 2 mL LB containing 0.2 mM IPTG. This cell suspension was incubated for 45 min at 37 °C. A part of the culture was plated on LB-agar containing 10 μg/mL tetracycline to measure the transformation efficiency of plasmid pVP146, and the rest was plated on LB containing 50 μg/mL ampicillin and 80 μg/mL X-gal to select for integrants (Amp^R^) and detect TLS events (*lacZ^+^* phenotype). Following the site-specific integration, the N^2^-furfuryl dG lesion is located either in the lagging strand (FBG151 derived strains) or in the leading strand (FBG152 derived strains). Cells were diluted and plated using the automatic serial diluter and plater EasySpiral Dilute (Interscience). Colonies were counted using the Scan 1200 automatic colony counter (Interscience). The integration rate was about 2,000 clones per picogram of a plasmid for a wild-type strain. Transformed cells were plated before the first cell division, and therefore, following the integration of the lesion-containing plasmid, blue-sectored colonies represent TLS events and pure white colonies represent damage avoidance (DA) events. The relative integration efficiencies of lesion-containing plasmids compared with their lesion-free equivalent, and normalized by the transformation efficiency of pVP146 plasmid in the same electroporation experiment, allow the overall rate of lesion tolerance to be measured.

### PALM Imaging

#### Sample preparation for microscopy and MMS treatment

Samples were prepared for microscopy following previously reported procedures(Thrall et al., 2017). In brief, glycerol stocks were streaked on LB plates containing 30 µg/mL kanamycin when appropriate. The day before imaging, a 3 mL “overday” LB culture was inoculated with a single colony and incubated for approximately 8 h at 37 °C on a roller drum. An overnight culture was prepared in 3 mL M9 medium supplemented with 0.4% glucose, 1 mM thiamine hydrochloride, 0.2% casamino acids, 2 mM MgSO_4_, 0.1 mM CaCl_2_, and 0.5 mM IPTG to induce expression of *ssb-mypet*. This overnight culture was inoculated with a 1:1,000 dilution of the overday culture and incubated overnight on a roller drum at 37 °C. On the day of imaging, 50 mL cultures in supplemented M9 medium were inoculated with a 1:200 dilution of the overnight culture and incubated at 37 °C shaking at 225 rpm.

Cells were harvested for imaging in early exponential phase when OD_600nm_ ≈ 0.15. Aliquots were removed and concentrated by centrifugation, then the concentrated cell suspension was deposited on an agarose pad sandwiched between cleaned coverslips. The coverslip in the optical path was cleaned by two 30 min cycles of sonication in ethanol and 1 M KOH and stored in deionized water. Agarose pads were prepared by heating 3% GTG agarose (NuSieve) in M9 medium supplemented with 0.4% glucose, 2 mM MgSO_4_, and 0.1 mM CaCl_2_, then casting a 500 µL volume of molten agarose between two cleaned 25 × 25 mm coverslips. For MMS treatment, MMS was added to the molten agarose at 100 mM immediately before casting the pad. Cells were incubated on the MMS agarose pad in a humidified chamber at RT for 20 min before imaging.

#### Microscopy

Imaging was performed on a customized Nikon TE2000 microscope described previously(Thrall et al., 2017). Briefly, 405 nm (Coherent OBIS, 100 mW) and 561 nm (Coherent Sapphire, 200 mW) laser excitation was used to activate and excite PAmCherry, and 514 nm (Coherent Sapphire, 150 mW) laser excitation was used to excite mYPet. Imaging was carried out with highly inclined thin illumination(Tokunaga et al., 2008), or near-TIRF illumination, by focusing the beams to the back focal plane (BFP) of a Nikon CFI Apo 100X/1.49 NA TIRF objective. The microscope was equipped with Chroma dichroic and emission filters (91032 Laser TIRF Cube containing a ZT405/514/561rpc dichroic filter, ZET442/514/561m emission filter, and ET525lp longpass filter). A Hamamatsu ImageEM C9100-13 EMCCD camera was used to record images with 250 ms exposure time. For all two-color PALM movies, the excitation sequence commenced with a high-power pre-bleaching period of 50 frames (∼ 120 W cm^−2^ 561 nm excitation) was followed by 10 frames of low-power pre-bleaching (∼ 12.5 W cm^−2^ 561 nm power). Then 10 frames were recorded to image SSB-mYPet (∼ 0.4 W cm^−2^ 514 nm power). Finally, PAmCherry fusions were activated and excited by continuous 405 nm (starting at ∼ 2.5 mW cm^−2^ power and increasing to ∼ 17.5 mW cm^−2^ power) and 561 nm (∼ 12.5 W cm^−2^ power) illumination. White light transillumination was used for acquisition of brightfield images of cells. All imaging experiments were repeated for at least three biological replicates (imaging cultures) on at least two different days.

#### Image analysis

As described previously(Thrall et al., 2017), image analysis was performed in MATLAB using MicrobeTracker(Sliusarenko et al., 2011) for cell segmentation, u-track for spot detection and tracking(Jaqaman et al., 2008), and custom code for other analysis. The point source detection algorithm(Aguet et al., 2013) in u-track was used to detect mYPet and PamCherry fusions and to fit them to 2D Gaussian point spread functions (PSFs). For PAmCherry fusions, a significance threshold of *α* = 10^−6^ was applied. Static tracks were identified by comparing the mean width of the PSF to the distribution in fixed cells; mobile molecules have broader PSFs due to motion blurring(Uphoff et al., 2013). Tracks with mean PSF width between 85.5 and 211.0 nm were determined to be static based on previous measurements(Thrall et al., 2017). A small number of cells containing PAmCherry localizations in the first PALM frame were excluded from analysis as a precaution against cross-talk from the mYPet channel. For SSB-mYPet, the first 5 frames of 514 nm excitation were averaged and analyzed with a significance threshold of *α* = 10^−5^. To remove a small number of poorly-fit foci, detected spots with a background level below the camera offset level (1,500 counts) were rejected. Other analysis parameters are as described previously(Thrall et al., 2017).

Colocalization analysis between SSB-mYPet foci and PamCherry fusions to Pol IV and RecQ^WH^-Pol IV^LF^ variants was performed as previously described(Thrall et al., 2017). In brief, the mean distance between each static Pol IV (or RecQ^WH^-Pol IV^LF^) track and the closest SSB focus was measured for each cell. These raw distances were aggregated across all cells and plotted. Additionally, radial distribution function analysis(Garza de Leon et al., 2017; Zawadzki et al., 2015) was used to determine the increased likelihood of localization at a distance *r* from the centroid of an SSB focus relative to random localization in the cell. For each cell, the raw Pol IV-SSB distances were measured; then random Pol IV-SSB distances were calculated by generating the same number of random Pol IV localizations within the same cell outline and keeping the same SSB centroid positions. After repeating this procedure for all cells, the experimental distance distribution was normalized by the random distribution, yielding the radial distribution function *g(r)*. This approach was repeated 100 times and the resulting *g(r)* curves were averaged to give the final result. Additionally, an independent random Pol IV-SSB distance distribution was generated and 100 random *g(r)* curves were averaged in the same way to give a mean random *g(r)* curve. Values of the experimental *g(r)* curves greater than 1 indicate enrichment relative to a random distribution, whereas deviations of the random *g(r)* curves from 1 indicate errors due to the finite sample size. Fig S4A-C illustrates this procedure and shows representative mean *g(r)* curves and replicates.

---

**Supplementary table 1.** *E. coli* strains and plasmids used in this study.

| Strains | Relevant genetic alterations | Genetic backgrounds | References or Sources |
| --- | --- | --- | --- |
| S6 | *Wild-type* | *MG1655* |  |
| S7 | Δ*dinB* | Δ*dinB::frt-kan-frt* | (Chang et al., 2019) |
| S185 | *ptet-dinB* | Δ*lamB::ptet-dinB-frt-kan-frt* | This study |
| S186 | *ptet-flag-dinB* | Δ*lamB::ptet-flag-dinB-frt-kan-frt* | This study |
| S195 | *ptet-flag-mypet* | Δ*lamB::ptet-flag-mypet-frt-kan-frt* | This study |
| S313 | *dinB*<sup>T120P</sup> | *dinB::dinB*<sup>T120P</sup>-*frt-kan-frt* | This study |
| S380 | *ptet-dinB* | Δ*dinB::frt-cm-frt* Δ*lamB::ptet-dinB-frt-kan-frt* | This study |
| S381 | *ptet-flag-dinB* | Δ*dinB::frt-cm-frt* Δ*lamB::ptet-flag-dinB-frt-kan-frt* | This study |
| S383 | *ptet-dinB*<sup>ΔC6</sup> | Δ*dinB::frt-cm-frt* Δ*lamB::ptet-dinB*<sup>ΔC6</sup>-*frt-kan-frt* | This study |
| S385 | *ptet-dinB*<sup>T120P</sup> | Δ*dinB::frt-cm-frt* Δ*lamB::ptet-dinB*<sup>T120P</sup>-*frt-kan-frt* | This study |
| S392 | *ptet-dinB*<sup>LF</sup> | Δ*lamB::ptet-dinB*<sup>LF</sup>-*frt-kan-frt* | This study |
| S398 | *ptet-flag-mypet* | Δ*dinB::frt-cm-frt* Δ*lamB::ptet-flag-mypet-frt-kan-frt* | This study |
| S490 | Δ*lafU* | Δ*lafU::frt-kan-frt* | This study |
| S502 | *dinB*<sup>Y79L</sup> | *dinB::dinB*<sup>Y79L</sup>-*frt-kan-frt* | This study |
| S519 | *sulAp-gfp-mut2* *dinB*<sup>lexA</sup> | Δ*attB::sulAp-gfp-mut2-frt-cm-frt* Δ*lafU::frt-kan-frt* *dinB*<sup>lexA</sup> | This study |
| S530 | *sulAp-gfp-mut2* | Δ*attB::sulAp-gfp-mut2-frt-cm-frt* Δ*lafU::frt-kan-frt* | (Chang et al., 2019) |
| S532 | *dnaQ*(e<sub>L</sub>) | Δ*dnaQ::dnaQ*(e<sub>L</sub>) Δ*yafT::frt-cm-frt* | (Chang et al., 2019) |
| S603 | *dinB*<sup>Y79L,T120P</sup> | *dinB::dinB*<sup>Y79L,T120</sup>-*frt-kan-frt* | This study |
| S611 | *dinB*<sup>T120A</sup> | *dinB::dinB*<sup>T120A</sup>-*frt-kan-frt* | This study |
| S612 | *dinB*<sup>T120S</sup> | *dinB::dinB*<sup>T120S</sup>-*frt-kan-frt* | This study |
| S613 | *dinB*<sup>T120V</sup> | *dinB::dinB*<sup>T120V</sup>-*frt-kan-frt* | This study |
| S614 | *dinB*<sup>T120D</sup> | *dinB::dinB*<sup>T120D</sup>-*frt-kan-frt* | This study |
| S644 | *dinB*<sup>T120P,Rim, ΔC6</sup>-*PAmCherry*<br>*ssb-mypet* | Δ*dinB::dinB*<sup>T120P,Rim, ΔC6</sup>-*PamCherry-frt-kan-frt*<br>Δ*lacZ::ssb-mypet-frt* *lexA51* | This study |
| S673 | *ptet-flag-polB* | Δ*lamB::ptet-flag-polB-frt-kan-frt* | This study |
| S685 | *ptet-flag-recQ* | Δ*lamB::ptet-flag-recQ-frt-kan-frt* | This study |
| S687 | *ptet-flag-recQ*<sup>WH</sup> | Δ*lamB::ptet-flag-recQ*<sup>WH</sup>-*frt-kan-frt* | This study |
| S742 | *sulAp-gfp-mut2* Δ*dinB* | Δ*attB::sulAp-gfp-mut2-frt-cm-frt* Δ*dinB::frt-kan-frt* | (Chang et al., 2019) |
| S800 | *dinB*<sup>T120P</sup> | Δ*lafU::frt-kan-frt* *dinB*<sup>T120P</sup> | This study |
| S802 | *ptet-dinB*<sup>1-230</sup> | Δ*dinB::frt-cm-frt* *ptet-dinB*<sup>1-230</sup>-*frt-kan-frt* | This study |
| S804 | *ptet-dinB*<sup>LF</sup> | Δ*dinB::frt-cm-frt* *ptet-dinB*<sup>LF</sup>-*frt-kan-frt* | This study |
| S852 | *sulAp-gfp-mut2* *dinB*<sup>T120P</sup> | Δ*attB::sulAp-gfp-mut2-frt-cm-frt* Δ*lafU::frt-kan-frt* *dinB*<sup>T120P</sup> | This study |
| S853 | *sulAp-gfp-mut2 dinB<sup>lexA</sup>*<br>*dinB<sup>T120P</sup>* | *ΔattB::sulAp-gfp-mut2-frt-cm-*<br>*frt ΔlafU::frt-kan-frt dinB<sup>lexA</sup>*<br>*dinB<sup>T120P</sup>* | This study |
| S902 | *ptet-flag-recQ<sup>WH</sup>-dinB<sup>LF</sup>* | *ΔlamB::ptet-flag-recQ<sup>WH</sup>-*<br>*dinB<sup>LF</sup>-frt-kan-frt* | This study |
| S929 | *recQ<sup>WH</sup>-dinB<sup>LF</sup>-PamCherry* | *ΔdinB::recQ<sup>WH</sup>-dinB<sup>LF</sup>-*<br>*PamCherry-frt-kan-frt* | This study |
| S930 | *recQ<sup>WH</sup>-dinB<sup>LF</sup>-PamCherry-flag*<br>*ssb-mypet* | *ΔdinB::recQ<sup>WH</sup>-dinB<sup>LF</sup>-*<br>*PamCherry-flag-frt-kan-frt*<br>*lacZ::ssb-mypet-frt lexA51* | This study |
| S937 | *ΔyafT ΔlafU pVP135* | *EVP22b ΔyafT::frt-cm-frt*<br>*ΔlafU::frt pVP135* | This study |
| S939 | *ΔyafT ΔlafU dinB<sup>T120P</sup> pVP135* | *EVP22b ΔyafT::frt-cm-frt*<br>*ΔlafU::frt dinB-T120P pVP135* | This study |
| S952 | *recQ<sup>WH,R503A</sup>-dinB<sup>LF</sup>-PamCherry* | *ΔdinB::recQ<sup>WH,R503A</sup>-dinB<sup>LF</sup>-*<br>*PamCherry-frt-kan-frt* | This study |
| S983 | *recQ<sup>WH(R503A)</sup>-dinB<sup>LF</sup> ssb-mypet* | *ΔdinB::recQ<sup>WH(R503A)</sup>-dinB<sup>LF</sup>-*<br>*PamCherry-frt-kan-frt*<br>*ΔlacZ::ssb-mypet-frt lexA51* | This study |
| S984 | *recQ<sup>WH</sup>-dinB<sup>LF(Rim, ΔC6)</sup>-*<br>*PamCherry ssb-mypet* | *ΔdinB::recQ<sup>WH</sup>-dinB<sup>LF(Rim, ΔC6)</sup>-*<br>*PamCherry-frt-kan-frt*<br>*ΔlacZ::ssb-mypet-frt lexA51* | This study |
| S1069 | *ptet-flag-recQ<sup>R448A</sup>* | *ΔlamB::ptet-flag-recQ<sup>R448A</sup>-frt-*<br>*kan-frt* | This study |
| S1071 | *ptet-flag-recQ<sup>R503A</sup>* | *ΔlamB::ptet-flag-recQ<sup>R503A</sup>-frt-*<br>*kan-frt* | This study |
| S1074 | *ptet-flag-exoI* | *ΔlamB::ptet-flag-exoI-frt-kan-*<br>*frt* | This study |
| S1075 | *ptet-flag-topoIII* | *ΔlamB::ptet-flag-topoIII-frt-*<br>*kan-frt* | This study |
| S1088 | *ptet-flag-exoI<sup>Q311A</sup>* | *ΔlamB::ptet-flag-exoI<sup>Q311A</sup>-frt-*<br>*kan-frt* | This study |
| S1089 | *ptet-flag-exoI<sup>R316A</sup>* | *ΔlamB::ptet-flag-exoI<sup>R316A</sup>-frt-*<br>*kan-frt* | This study |
| S1090 | *ptet-flag-exoI<sup>R414A</sup>* | *ΔlamB::ptet-flag-exoI<sup>R414A</sup>-frt-*<br>*kan-frt* | This study |
| S1091 | *ptet-flag-polA* | *ΔlamB::ptet-flag-polA-frt-kan-*<br>*frt* | This study |
| S1093 | *polA-recQ<sup>WH</sup>* | *ΔpolA::polA-recQ<sup>WH</sup>-frt-kan-frt* | This study |
| S1094 | *polA-recQ<sup>WH(R425A,R503A)</sup>* | *ΔpolA::polA-*<br>*recQ<sup>WH(R425A,R503A)</sup>-frt-kan-frt* | This study |
| S1095 | *polA<sup>+</sup>* | *ΔpolA::polA-frt-kan-frt* | This study |
| S1158 | *recQ<sup>WH</sup>-dinB<sup>LF(Rim, ΔC6)</sup>-*<br>*PamCherry-flag ssb-mypet* | *ΔdinB::recQ<sup>WH</sup>-dinB<sup>LF(Rim, ΔC6)</sup>-*<br>*PamCherry-flag-frt-kan-frt*<br>*lacZ::ssb-mypet-frt lexA51* | This study |
| S1162 | *recQ<sup>WH(R503A)</sup>-dinB<sup>LF</sup>-*<br>*PamCherry-flag ssb-mypet* | *ΔdinB::recQ<sup>WH(R503A)</sup>-dinB<sup>LF</sup>-*<br>*PamCherry-flag-frt-kan-frt*<br>*lacZ::ssb-mypet-frt lexA51* | This study |
| S1168 | *dinB<sup>ΔC6</sup>* | *ΔlafU::frt-kan-frt*<br>*ΔdinB::dinB<sup>ΔC6</sup>* | This study |
| S1171 | *dinB<sup>T120P, ΔC6</sup>* | *ΔlafU::frt-kan-frt*<br>*ΔdinB::dinB<sup>T120P, ΔC6</sup>* | This study |
| S1177 | *ptet-flag-polA-recQ<sup>WH</sup>* | *ΔlamB::ptet-flag-polA-*<br>*recQ<sup>WH</sup>-frt-kan-frt* | This study |
| S1178 | *ptet-flag-polA-recQ<sup>WH(R425A,R503A)</sup>* | *ΔlamB::ptet-flag-polA-recQ<sup>WH(R425A,R503A)</sup>-frt-kan-frt* | This study |
| S1179 | *ptet-flag-recQ<sup>WH</sup>-dinB<sup>LF(ΔC6)</sup>* | *ΔlamB::ptet-flag-recQ<sup>WH</sup>-dinB<sup>LF(ΔC6)</sup>-frt-kan-frt* | This study |
| S1205 | *dnaQ(e<sub>L</sub>) dinB<sup>T120P</sup>* | *ΔdnaQ::dnaQ(e<sub>L</sub>) ΔyafT::frt-cm-frt ΔlafU::frt-kan-frt dinB<sup>T120P</sup>* | This study |
| S1252 | *hda-recQ<sup>WH</sup>* | *Δhda::hda-recQ<sup>WH</sup>-frt-kan-fr* | This study |
| S1253 | *hda-recQ<sup>WH(R425A,R503A)</sup>* | *Δhda::hda-recQ<sup>WH(R425A,R503A)</sup>-frt-kan-fr* | This study |
| S1254 | *hda<sup>+</sup>* | *Δhda::hda-frt-kan-fr* | This study |
| S1295 | *ptet-crfc* | *ΔlamB::ptet-crfc-frt-kan-frt* | This study |
| S1317 | *ptet-crfc-recQ<sup>WH</sup>* | *ΔlamB::ptet-crfc-recQ<sup>WH</sup>-frt-kan-frt* | This study |
| S1318 | *ptet-crfc-recQ<sup>WH(R425A,R503A)</sup>* | *ΔlamB::ptet-crfc-recQ<sup>WH(R425A,R503A)</sup>-frt-kan-frt* | This study |
| EVP22b | *ΔuvrA ΔmutS pVP135* | *FBG151 ΔuvrA::frt ΔmutS::frt pVP135* | (Esnault et al., 2007; Pagès et al., 2012) |
| EVP23b | *ΔuvrA ΔmutS pVP135* | *FBG152 ΔuvrA::frt ΔΔmutS::frt pVP135* | (Esnault et al., 2007; Pagès et al., 2012) |
| EVP886 | *ΔdinB pVP135* | *EVP22b ΔdinB::frt pVP135* | This study |
| EVP887 | *ΔdinB pVP135* | *EVP23b ΔdinB::frt pVP135* | This study |
| S1106 | *ΔlafU pVP135* | *EVP22b ΔlafU::FRT pVP135* | This study |
| S1107 | *dinB<sup>T120P</sup> pVP135* | *EVP22b ΔlafU::FRT dinB<sup>T120P</sup> pVP135* | This study |
| EVP957 | *dinB<sup>T120P</sup> pVP135* | *EVP23b ΔlafU::FRT dinB<sup>T120P</sup> pVP135* | This study |
| EVP392 | *ΔumuDC ΔpolB ΔdinB pVP135* | *EVP22b ΔumuDC::frt polB::frt dinB::frt pVP135* | This study |
| EVP415 | *ΔumuDC ΔpolB ΔdinB pVP135* | *EVP23b ΔumuDC::frt ΔpolB::frt ΔdinB::frt pVP135* | This study |
| JEK687 | *ptet-flag-mypet* | *ΔlamB::ptet-flag-mypet-frt-kan-frt* | This study |
| JEK762 | *ssb-mypet* | *ΔlacZ::ssb-mypet-frt lexA51* | (Thrall et al., 2017) |
| JEK766 | *dinB-PAmCherry ssb-mypet* | *ΔdinB::dinB-PAmCherry-frt-kan-frt ΔlacZ::ssb-mypet-frt lexA51* | (Thrall et al., 2017) |
| JEK781 | *dinB-PAmCherry-flag ssb-mypet* | *ΔdinB::dinB-PAmCherry-flag-frt-kan-frt ΔlacZ::ssb-mypet-frt lexA51* | (Thrall et al., 2017) |
| JEK790 | *dinB<sup>Rim,ΔC6</sup>-PAmCherry ssb-mypet* | *ΔdinB::dinB<sup>Rim,ΔC6</sup>-PAmCherry-frt-kan-frt ΔlacZ::ssb-mypet-frt lexA51* | (Thrall et al., 2017) |
| ET215 | *dinB<sup>T120P</sup>-PAmCherry* | *ΔdinB::dinB<sup>T120P</sup>-PAmCherry-frt-kan-frt* | This study |
| ET216 | *dinB<sup>T120P</sup>-PAmCherry-flag* | *ΔdinB::dinB<sup>T120P</sup>-PAmCherry-flag-frt-kan-frt* | This study |
| ET227 | *dinB<sup>T120P</sup>-PAmCherry ssb-mypet* | *dinB::dinB<sup>T120P</sup>-PAmCherry-frt-kan-frt ΔlacZ::ssb-mypet-frt lexA51* | This study |
| ET230 | *dinB<sup>T120P</sup>-PAmCherry-flag ssb-mypet* | *dinB::dinB<sup>T120P</sup>-PAmCherry-flag-frt-kan-frt ΔlacZ::ssb-mypet-frt lexA51* | This study |

| Plasmids | Relevant inserts | References or Sources |
| --- | --- | --- |
| pVP135 | Expression of phage lambda integrase/excisionase | (Pagès et al., 2012) |
| pVP146 | Internal control for transformation efficiency | (Pagès et al., 2012) |
| pAP1 | Parental plasmid for construction of N<sup>2</sup>-furfuryl dG-containing plasmid | This study |
| pGP22 | Parental plasmid for construction of N<sup>2</sup>-furfuryl dG-containing plasmid | This study |

